# Critical role for astrocyte NAD^+^ glycohydrolase in myelin injury and regeneration

**DOI:** 10.1101/2020.06.10.143941

**Authors:** Monica R. Langley, Chan-Il Choi, Thais R. Peclat, Yong Guo, Whitney Simon, Hyesook Yoon, Laurel Kleppe, Claudia F. Lucchinetti, Claudia C.S. Chini, Eduardo N. Chini, Isobel A. Scarisbrick

## Abstract

Western-style diets cause disruptions in myelinating cells and astrocytes within the mouse CNS. CD38 has increased expression in the cuprizone and EAE demyelination models and is the main NAD^+^ depleting enzyme in CNS tissue. Altered NAD^+^ metabolism has been linked to both high fat consumption and Multiple Sclerosis (MS). We identified increased CD38 expression in the male mouse spinal cord following chronic high fat consumption or focal lysolecithin-induced demyelinating injury as well as in reactive astrocytes within an active MS lesion. CD38-catalytically inactive mice are significantly protected from high fat-induced NAD^+^ depletion, oligodendrocyte loss, oxidative damage, and astrogliosis. 78c, a CD38 inhibitor, increased NAD^+^ and attenuated neuroinflammatory changes in astrocytes induced by saturated fat. Conditioned media from saturated fat-treated astrocytes impaired oligodendrocyte differentiation pointing to indirect mechanisms of oligodendrogliopathy. Combined saturated fat and lysolecithin demyelination in cerebellar slices resulted in additional deficits in myelin proteins that were mitigated by concomitant 78c treatment. Importantly, oral 78c increased counts of oligodendrocytes and remyelinated axons after focal demyelination. Our findings suggest high fat diet impairs oligodendrocyte survival and differentiation through astrocyte-linked mechanisms mediated by the NAD^+^ase CD38, and highlight the use of CD38 inhibitors as potential therapeutic candidates to improve myelin regeneration.

## Introduction

Obesity is a key risk factor for developing multiple sclerosis (MS)(1, 2). Moreover, recent studies suggest that obesity and related metabolic comorbidities also influence disease progression and disability burden in individuals affected by MS (3). Although obesity-induced white matter abnormalities and loss of oligodendrocytes within the central nervous system (CNS) following chronic consumption of high fat diets (HFD) has been found in rodents, how diet influences white matter integrity in humans and interacts with MS pathophysiology is largely unknown (4–6).

Astrocyte activation occurs following chronic HFD feeding in the CNS (7, 8) and can exert oligotoxic effects (9). Building and maintenance of myelin membranes requires oligodendrocyte fatty acid synthesis, yet oligodendrocytes can utilize lipids from the diet or synthesized by astrocytes (10). The potential to target astrocytes to improve outcomes in MS is already suggested since some currently prescribed MS disease modifying therapies impact astrocytes (11). Prior studies regarding the role of HFDs in demyelination models have utilized immune-mediated models such as experimental autoimmune encephalomyelitis (EAE) and demonstrate exacerbation of peripheral immune responses or neuroinflammation with HFD consumption (12–15). There are few, if any, studies regarding the effects of HFD on the capacity for remyelination. Since oligodendrocyte function is intimately linked to neighboring glial cells and neuronal activity, in addition to microenvironmental factors, the most promising interventions will likely involve a combinatorial approach.

CD38 is an NAD^+^-degrading enzyme expressed by astrocytes and was recently shown to be upregulated in peripheral immune- and toxin-mediated demyelination models (15–18). CD38 is the main NAD^+^ase in many mammalian tissues and regulates tissue and cellular NAD^+^ homeostasis as well as NAD^+^ dependent enzymes such as PARPs and Sirtuins (19, 20). Global CD38 knockout confers protection against demyelination, T cell responses, and EAE disease severity as well as HFD-induced metabolic dysfunction in rodent models (15, 18, 21, 22). Here we test the hypotheses that HFD consumption promotes oligodendrocyte loss in a CD38-dependent manner and that CD38 inhibition will enhance myelin regeneration. First, we established increased CD38 expression following HFD consumption. Our previous studies indicate that high fat diets lead to a reduction in oligodendrocytes and their progenitors within the mouse CNS (4, 6, 8). Next, we demonstrated that CD38 catalytically inactive mice fed a HFD were protected from HFD-mediated loss of oligodendrocytes with concomitant increases in NAD^+^ levels and reduced neuroinflammation and oxidative damage. *In vitro* approaches identified oligodendrocyte direct and astrocyte-linked indirect mechanisms by which excessive fatty acid impairs myelinating cell function. Finally, the value of a small molecule inhibitor of CD38, 78c, was then identified as a novel protective strategy to enhance myelin regeneration in a model of focal demyelinating injury. Together these findings highlight a novel indirect pathway of myelin injury involving pathogenic changes in astrocytes. Moreover, these changes can be attenuated by inhibition of CD38 in the context of chronic HFD consumption and targeted to improve myelin regeneration in the CNS.

## Results

### Astrocytes increase CD38 in the spinal cord following chronic HFD or demyelination

Our recent studies revealed spinal cord changes involving ER stress, mitochondrial dysfunction, and oxidative stress in the spinal cords of mice chronically consuming a HFD and the direct potential of saturated fat palmitate (PA) to inhibit differentiation of primary murine oligodendrocytes from their progenitors or neural stem cells (4, 6). Although saturated fat inhibited markers of differentiation *in vitro*, it was not oligotoxic, suggesting a more complex mechanism at play *in vivo* (4). Considering the emerging role of astrocytes in the pathology of various neurological disorders (9, 11, 23) and in the CNS following chronic HFD (7, 8, 24), along with their known essential metabolic and functional links with oligodendrocytes (25, 26), we decided to address the possibility of indirect effects of HFD mediated by astrocytes (Figure 1A).

**Figure 1.**
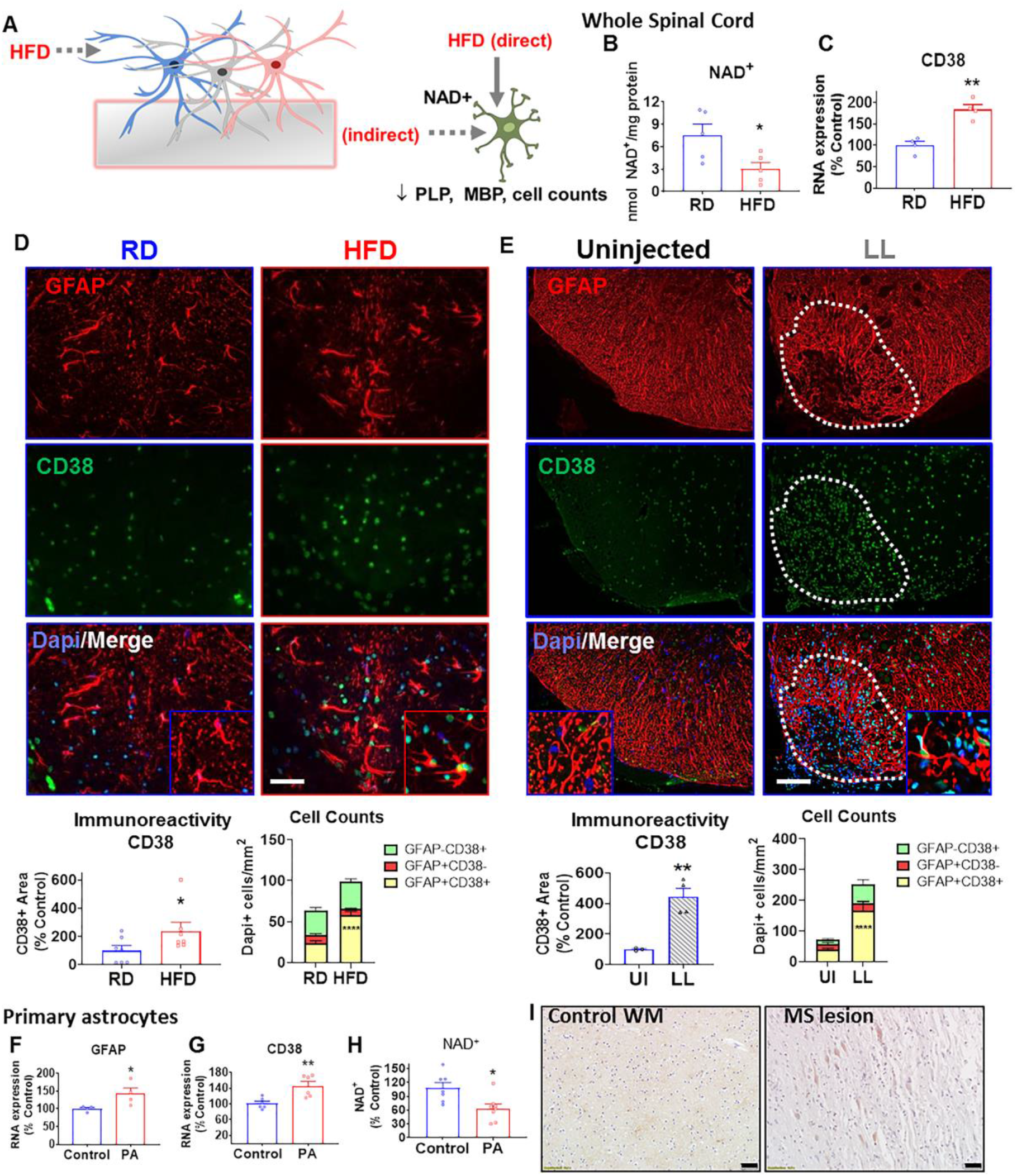
CD38 expression in astrocytes of chronic HFD-fed mice and in a model of demyelination. **A** Schematic representation of previously shown direct effects of high fat diet (HFD) on oligodendrocyte lineage cells including apoptosis and impaired differentiation with potential astrocyte-mediated indirect effects, represented by dotted lines, that comprise the hypothesis to be tested. **B** Spinal cord tissue of regular diet (RD) and high fat diet (HFD) mice showing a depletion of NAD^+^ after 12 wk (n=5 mice/group) as determined by cycling assay. **C** Increased RNA expression of CD38 (n=4), assessed by qPCR, in spinal cord tissue of HFD-fed mice when compared to RD-fed mice. **D-E** Immunofluorescence staining with double-labeling and corresponding quantification of immunoreactivity and cell counts indicates increased CD38 (green) immunoreactivity in astrocytes (GFAP, red) within the spinal cords of HFD-fed mice (***D***, n=7) or in the lysolecithin (LL) model of experimental demyelination (***E***, n=4)versus the uninjected (UI) side. Broken white line designates lesion boundary. Scale bar = 50 µm (***D***) and 100µm (***E***). **F-H** High fat consumption was modeled in primary murine astrocyte cultures by including saturated fat palmitate in the media (PA, 100 µM, 24 h). Increases in RNA expression of GFAP (***F,*** n=4) and CD38 (**G,** n=6) were observed in PA-containing astrocyte cultures. ***H***, Depleted NAD^+^ levels were found in PA-treated astrocytes when assessed using an NAD^+^ cycling assay, n=7-8. **I** Representative microscopic images of CD38 staining in control brain tissue and an MS lesion with active demyelinating activity. Scale bar = 50 µm. Bar graphs represent the mean +/− SEM. Asterisks indicate levels of statistical significance as determined by Student’s t-test, where *, p<0.05; **, p<0.01; ***, p<0.001.

NAD^+^ is a key mediator of cellular energy status and is involved in mitochondrial function and quality control processes primarily through the activity of sirtuins that require NAD^+^ as a cofactor (19, 20, 27). Exercise is known to increase NAD^+^ levels in various tissues (22), and protect against “Western-style diet”-induced deficits in myelin and myelinating cells in two recent studies (5, 6). Our recent untargeted metabolomics profiling of spinal cord tissue identified NAD^+^ metabolism as one of the top pathways impacted by HFD consumption (4), which prompted us to quantify spinal cord NAD^+^ levels using a cycling assay. Spinal cord NAD^+^ levels from HFD-fed mice were less than half that of RD-fed mice (p=0.029, t=2.639, df=8; Figure 1B).

Since CD38 is the main NAD^+^-glycohydrolase in many mammalian tissues(19, 20, 27), we next quantified CD38 expression in spinal cord astrocytes to establish a potential link. Quantification by qPCR indicated higher RNA expression of CD38 in HFD-fed mice compared to regular diet-fed controls (p=0.001, t=5.626, df=6; Figure 1 C). Consistent with findings demonstrating CD38 is primarily localized to astrocytes within the CNS (16–18, 28), increases in CD38 immunoreactivity were observed in GFAP^+^ astrocytes following HFD (p=0.038, t=1.937, df=12; Figure 1D). The HFD-mediated effects mirror the increased CD38 expression following lysolecithin-induced demyelination shown here (p=0.004, t=5.105, df=6; Figure 1E) and in EAE (15) and cuprizone (18) associated demyelination studies. Similar to the HFD spinal cords, primary murine astrocytes cultured with PA (100 µM, 24 h) showed increased GFAP (p=0.046, t=2.507, df=6; Figure 1F) and CD38 (p=0.005, t=3.576, df=10; Figure 1G) RNA expression and depleted NAD^+^ levels (p=0.011, t=2.962, df=13; Figure 1H). Collectively, these data suggest that HFD can induce astrocytosis in the spinal cord or cultures, which increases expression of CD38 and subsequently depletes NAD^+^ levels.

Microscopic imaging of human brain sections demonstrated extensive CD38 immunoreactivity in white matter astrocytes within control brain tissue and increased CD38 in some hypertrophic reactive astrocytes within MS lesion tissue with active demyelinating activity (Figure 1I). The expression and significance of CD38 immunoreactivity among various astrocyte subtypes and disease phenotypes needs to be further characterized, but is likely not specific to MS since GFAP+CD38+ cells were found to be increased in HIV-1-associated encephalopathy (HIVE) brain tissue sections (29).

### CD38 catalytically inactive (CD38ci) mice are resistant to HFD-induced oligodendrocyte loss

To determine the potential therapeutic significance of CD38 upregulation to HFD-induced oligodendrocyte loss, we collected spinal cords from wildtype (WT) and CD38-catalytically inactive (CD38ci) mice fed a regular diet (RD) or HFD for 30 wk beginning at 50 wk of age for histological analysis (Figure 2A). CD38ci mice have the CD38 gene knocked out and reinserted with a mutation at E230 of the CD38 gene, rendering the CD38-NAD^+^ase catalytic site inactive, while preserving overall expression levels of CD38 (30). To confirm that preservation of NAD^+^ levels was achieved in the CD38ci mice, we next measured NAD^+^ levels on spinal cord tissue from the same cohort of mice. As expected based on our data in Figure 1, NAD^+^ levels were significantly reduced following chronic HFD consumption in wildtype mice (F(_1, 24_)=1.785, p=0.004; Figure 2B). Moreover, in CD38ci mice, which had diminished CD38 activity (F(_1, 16_)=75.13, p<0.0001; Figure 2C), NAD^+^ levels were significantly higher overall (F(_1, 24_)=33.90, p<0.0001) and in the CD38ci-HFD mice compared to the wildtype-HFD mice (p<0.0001). As seen in our previous studies (4, 6), there was a significant reduction in the number of PDGFRα^+^ (F(_1, 16_)= 16.61, p=0.0003 dorsal column and F(_1, 16_)= 17.77, p=0.0007 ventral; Figure 2E) and Olig2^+^ oligodendrocyte progenitors (F(_1, 26_)= 9.066, p=0.006 dorsal column and F(_1, 26_)= 24.90, p<0.0001 ventral; Figure 2 E) as well as GST3^+^ mature oligodendrocytes (F(_1, 22_)= 14.53, p=0.001 dorsal column and F_(1, 25)_= 10.23, p=0.004 ventral; Figure 2F) in the HFD-fed wildtype mice compared to regular diet-fed mice. However, CD38ci mice fed HFD did not have reductions in oligodendrocyte lineage cell counts compared to their regular diet genotype controls (p=0.682), yet had significantly higher PDGFRα^+^ (p=0.0397 dorsal column and p=0.0198 ventral white matter) and GST3^+^ (p=0.0001 dorsal column) cell counts versus WT-HFD mice. The improved cell numbers are likely due to decreased apoptosis rather than proliferative changes, as evidenced by decreased cleaved-caspase-3 staining (F_(1, 16)_= 5.48, p=0.0224 ventral; Figure 2E) in the CD38ci-HFD versus WT-HFD group and no significant difference in Ki67. Across spinal cord regions, no significant changes in MBP due to genotype or diet group were found (F_(1, 28)_= 0.002, p= 0.966, and F_(1, 28)_= 0.683, p=0.415, respectively, Figure 2D). Overall, these data suggest that oligodendrocyte lineage cell survival is preserved in CD38ci mice following chronic HFD consumption.

**Figure 2.**
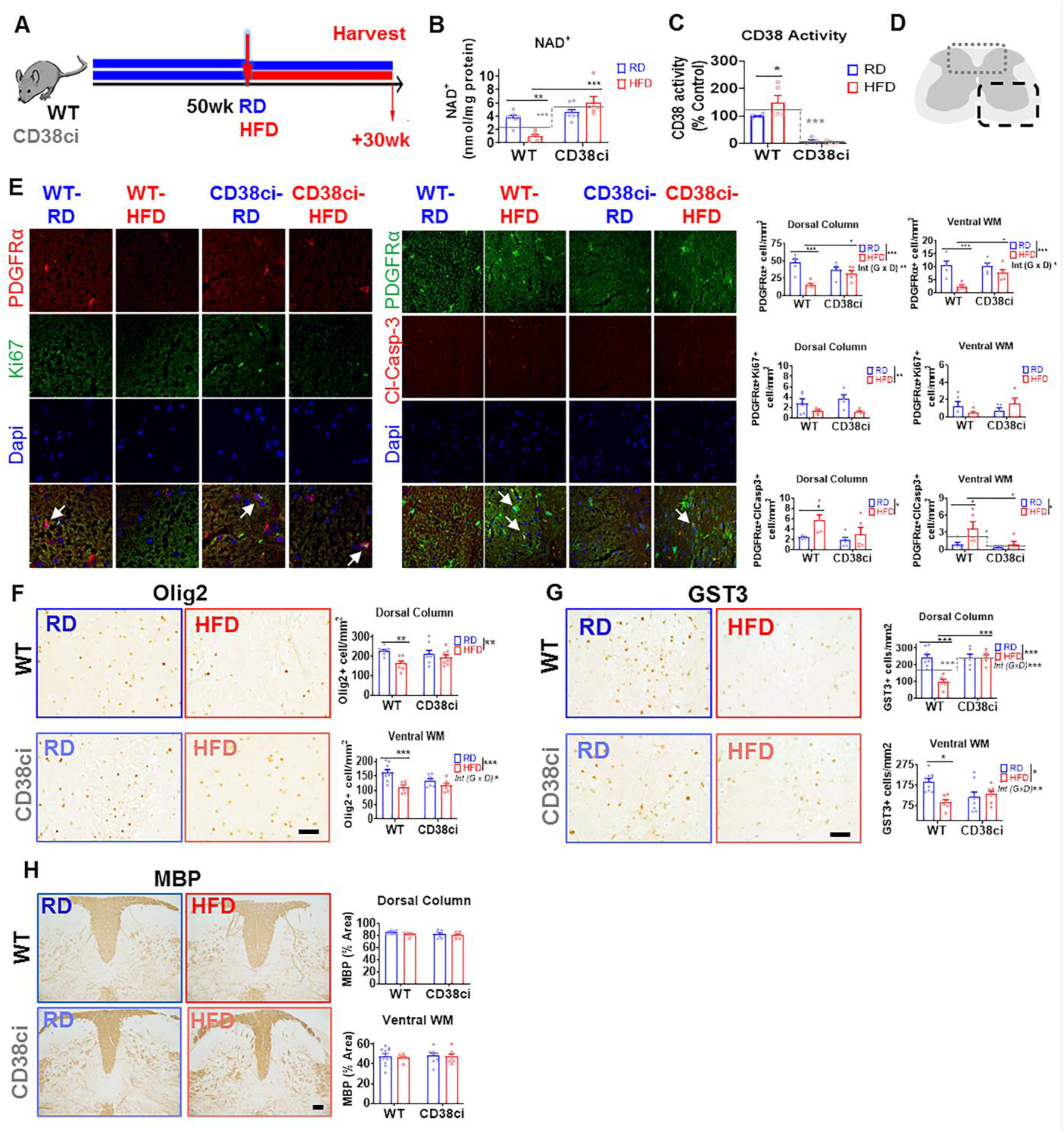
CD38-catalytically inactive (CD38ci) mice are resistant to HFD-induced oligodendrocyte loss. Histological analysis from wildtype (WT) and CD38-catalytically inactive mice (CD38ci) fed a regular diet (RD) or high fat diet (HFD) for 30 wk beginning at 50 wk of age. **A** Schematic depicting experimental design. **B** NAD^+^ levels as determined by cycling assay indicate reductions in WT-HFD mice compared to WT-RD and increased NAD^+^ levels in the CD38ci mice versus WT. **C** CD38 activity levels by etheno-NAD^+^ assay show impaired activity in CD38ci mice versus WTs and higher activity in WT-HFD mice than WT-RD (n=5/group). **D**, Outlined regions depict regions imaged, including dorsal column (broken gray line) and ventral white matter (broken black line). **E** PDGFRα double immunofluorescent staining with Ki67 or cleaved-caspase-3 shows a depletion of oligodendrocyte progenitor cells with HFD that was significantly attenuated in CD38ci mice versus wildtype. Increased cleaved-caspase-3 and PDGFRα co-labeled cells were found in HFD versus RD WT mice, while fewer cleaved-caspase-3 and PDGFRα co-labeled cells we observed in the CD38ci HFD than WT HFD ventral WM. **F-G** Olig2^+^ oligodendrocyte lineage cells (***F***) and GST3^+^ mature oligodendrocytes (***G***) were significantly depleted in HFD-fed WT mice, yet CD38ci mice fed a HFD had no reduction in oligodendrocyte numbers. When compared to WT-HFD mice, CD38ci-HFD mice had significantly more oligodendrocytes remaining in the dorsal column (representative images pictured). Scale bar = 50 µm. Gray lines indicate genotype means, while gray broken lines indicate comparison of overall genotype effect. **H** Representative images of immunohistochemical staining confirming no significant changes in MBP immunoreactivity across genotype or diet groups. Scale bar= 100 µm. Bar graphs represent mean +/− SEM. Two-way ANOVA followed by Bonferroni post-hoc test (n=7-10 mice/group). Asterisks indicate levels of statistical significance when compared to control, where *, p<0.05; **, p<0.01; ***, p<0.001.

### Diminished neuroinflammatory responses and oxidative damage in CD38ci mice following chronic HFD

Neuroinflammation, including complex interplay between microglia and astrocytes, is now appreciated to contribute to oligotoxicity and MS pathophysiology (9, 25). To explore the potential impact of blocking CD38 catalytic activity on CNS neuroinflammation following chronic HFD consumption, we next performed immunohistochemical analysis on the spinal cords from wildtype and CD38ci mice fed regular or HFD for markers of microglia and astrocytes. Quantification of the thresholded area considered positive for ionized calcium-binding adapter molecule 1 (IBA1) demonstrated decreased microglia numbers or activity (F_(1, 21)_= 5.079, p=0.030; Figure 3A) in the ventral grey matter of spinal cords from CD38ci mice compared to wildtype mice overall. In addition, HFD-fed CD38ci mice showed significantly reduced IBA1 immunoreactivity in the ventral grey and white matter of the spinal cord compared to wildtype mice fed a HFD. Although HFD did not significantly increase IBA1-immunoreactivity in this study, this is consistent with a recent Western-style diet study from our laboratory and older mice may already have residually higher levels (8, 31). Considering that astrocytes have endogenous expression of CD38 as well as elevated CD38 expression following HFD or demyelination (16–18), we also investigated astrocyte reactivity in these mice. In the ventral spinal cord, quantification of astrocyte marker glial fibrillary acidic protein (GFAP, F_(1, 28)_= 12.66, p=0.0014; Figure 3B) demonstrated significantly reduced positive area in the white matter of CD38ci mice compared to wildtype mice. Wildtype HFD-fed mice showed significantly more astrocyte reactivity compared to either regular diet (p=0.023) or HFD-fed CD38ci mice (p=0.043).

**Figure 3.**
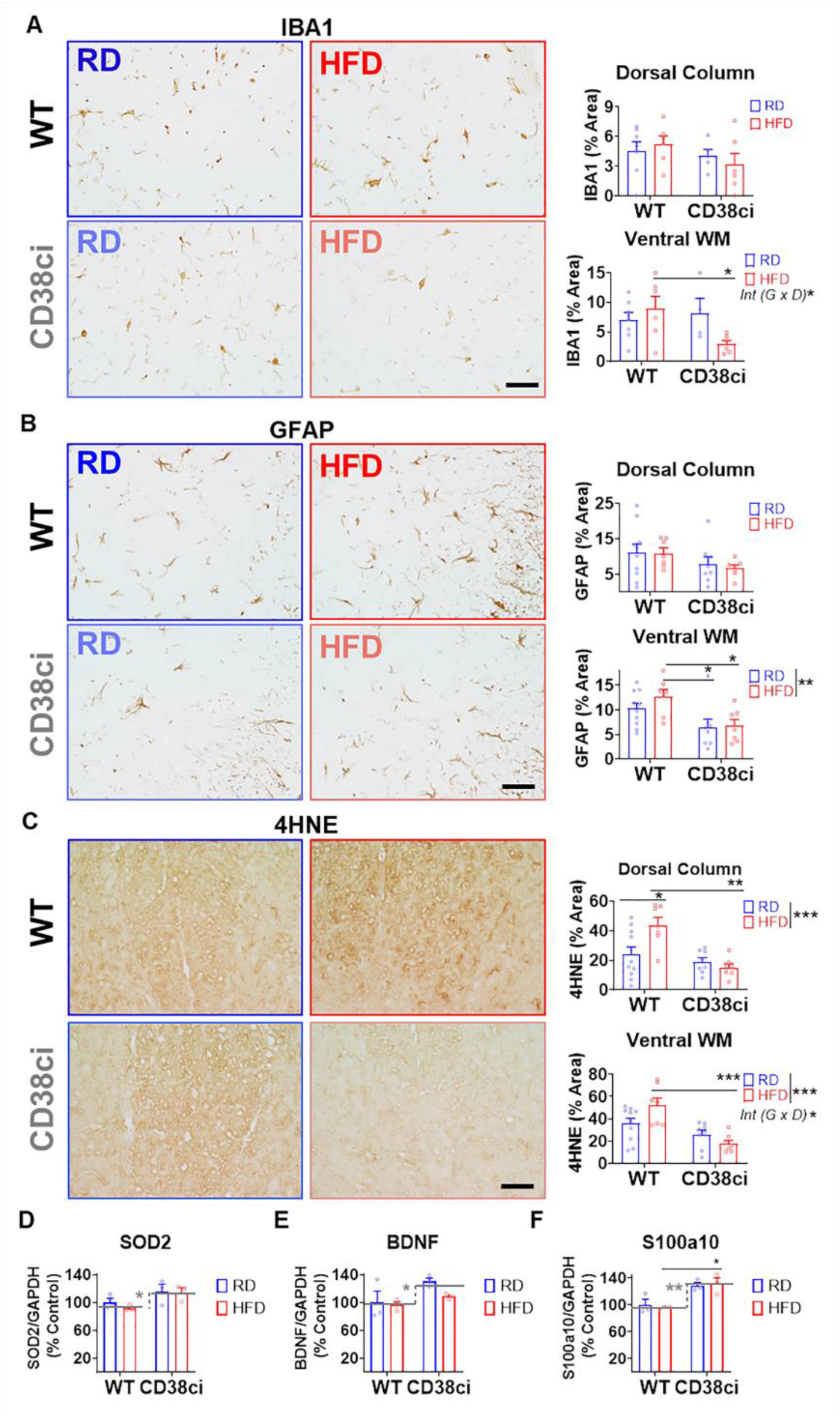
Diminished neuroinflammatory responses in CD38 catalytically inactive (CD38ci) mice following chronic HFD. Histological analysis from wildtype (WT) and CD38-catalytically inactive mice (CD38ci) fed a regular diet (RD) or high fat diet (HFD) for 30 wk beginning at 50 wk of age. Scale bar= 50 µm. and n=7-10 mice/group. **A** Representative images of the ventral white matter and immunohistochemical analysis of spinal cord regions demonstrates decreased microgliosis (IBA1) in CD38ci mice. When compared to WT-HFD mice, CD38ci-HFD mice had significantly less IBA1^+^ immunoreactivity in the spinal cord ventral gray and white matter. **B** Representative images of the ventral white matter and corresponding quantification confirm reduced astrogliosis (GFAP) in the spinal cords of CD38 catalytically inactive (CD38ci) mice overall when compared to wildtype (WT) mice. Decreased GFAP^+^ immunoreactivity in the ventral white matter when compared to HFD mice. **C** Representative images of the dorsal column and quantification of the spinal cords confirming significant changes in 4HNE immunoreactivity, an oxidative stress marker, between genotypes for HFD mice. **D-F** qRT-PCRs data indicting differences in antioxidant (F) and pro-repair astrocyte marker genes (G-H, BDNF and S100a10, respectively) in the CD38ci mice versus controls. Bar graphs represent mean +/− SEM. Two-way ANOVA with Bonferroni post-test. Asterisks indicate levels of statistical significance, where *, p<0.05 and **, p<0.01. Gray lines on graphs indicate genotype means, while gray broken lines with asterisks indicate a significant effect of genotype overall.

Given our previous data suggesting increased oxidative damage markers in the CNS following chronic consumption of HFDs (4, 6) and the significance of oxidative stress in glia of MS lesions (32), we next sought to compare the extent of oxidative damage in the spinal cords of wildtype and CD38ci mice fed regular or HFD. More abundant lipid peroxidation marker 4-hydroxynonenal (4HNE) was present in the HFD-fed wildtype mice than regular diet-fed wildtype mice in the spinal cord dorsal column (F_(1, 28)_= 3.358, p=0.019, Figure 3C). Importantly, a significant reduction in 4HNE was found in the HFD-CD38ci mice compared to HFD-wildtype mice in the dorsal column and ventral spinal cord (grey and white matter). Superoxide dismutase 2 (SOD2) is a mitochondrial antioxidant that can protect from high fat-induced changes in oxidative stress and metabolism (33), while brain-derived neurotrophic factor (BDNF) is permissive to myelination (23) and S100a10 is a recently defined pro-repair astrocyte maker (9). Next, our histological findings were further substantiated by observed gene expression increases in SOD2 (F_(1, 8)_= 7.721, p=0.024; Figure 3D) and pro-repair astrocyte markers genes BDNF(F_(1, 8)_= 5.685, p= 0.044; Figure 3E) and S100a10 (F_(1, 8)_= 24.73, p= 0.0011; Figure 3 F) in the CD38ci mice, as determined by qRT-PCR. Overall, the observed reductions in neuroinflammation and oxidative stress markers and increased NAD^+^ levels in the CD38ci mice likely provide a more suitable microenvironment for oligodendrocyte survival, differentiation, and repair following chronic HFD consumption.

### CD38 inhibition with 78c improves myelin repair following lysolecithin-induced focal demyelination of the ventral spinal cord

78c is a thiazoloquin(az)olin(on)e CD38 inhibitor that increases NAD^+^ levels in various tissues and cell types and is orally bioavailable (30, 34). To determine whether pharmacologic inhibition of CD38 affects the potential for remyelination, we administered 78c to mice in each diet group (regular or high fat) and used histological approaches to investigate numbers of oligodendrocyte lineage cells and remyelinated axons in the lysolecithin model of demyelination. Six-week old male wildtype mice were given either a regular or HFD for 5 weeks, then split into additional groups: receiving the same diet with or without 78c (600ppm) (Figure 4A). All mice were subjected to a focal demyelinating injury to the ventral spinal cord by microinjection of 1% lysolecithin.

**Figure 4.**
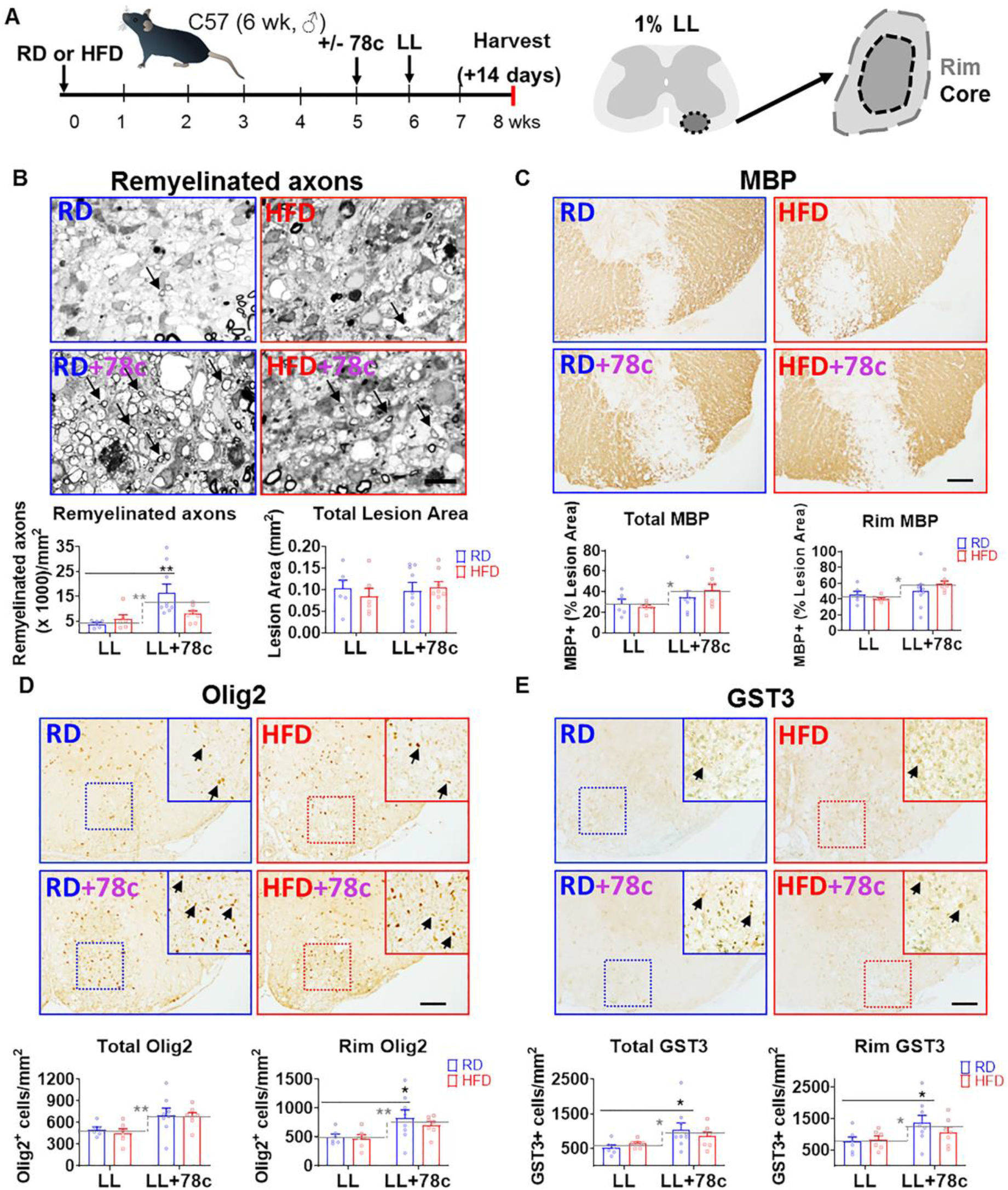
CD38 inhibition with 78c improves myelin repair following lysolecithin-induced focal demyelination to the ventral spinal cord. Eight week old male C57Bl6 mice were fed a regular diet (RD) or high fat diet (HFD) for 5 wks, then split into two additional groups: one receiving the same diet and the other half receiving the respective diet with 78c added (600 ppm). One week later, a focal demyelinating lesion to the ventral spinal cord was induced by lysolecithin (LL, 1% with Evans Blue dye) and mice were allowed to recover for two weeks. Lesion boundary regions were determined by limiting to the area of continuous myelin loss (core) or by including the entire inflammatory lesion (rim) based on immunohistochemical analyses. **A** Schematic depicting experimental design and lesion boundaries. **B** Representative images and corresponding analysis illustrating the mean number of remyelinated axons after focal lysolecithin-mediated demyelination of the ventral spinal cord was increased in 78c-treated mice at 14 days post injury when compared to LL only mice (**, p<0.01, two-way ANOVA with Bonferroni post-test) as assessed by paraphenylenediamine (PPD)-stained semi-thin sections of araldite-embedded spinal cord tissue. Scale bar= 20 µm. **C** Immunohistochemical staining of the total lesion and lesion rim identified significant changes in MBP immunoreactivity in 78c treated animals. **D-E** Olig2^+^ oligodendrocyte lineage cells (OLCs, *D*) and GST3^+^ mature oligodendrocytes (*E*) were significantly higher in 78c-RD mice when compared to RD controls. Bar graphs represent mean +/− SEM. Two-way ANOVA (n=7-10 mice/group). *, p<0.05 and **, p<0.01. Gray lines on graphs indicate genotype means, while gray broken lines indicate a significant effect of genotype overall.

After two weeks of recovery, lesions were collected from perfused mice and used for subsequent counting of remyelinated axons and myelinating cells. At the study endpoint, HFD mice had gained significantly more weight than their RD counterparts (F_(1, 36)_=51.91, p<0.0001, Supplemental Figure 1A), irrespective of genotype. While 78c treatment did not have a significant effect on body weights, mice given 78c consumed less food (Supplemental Figure 1B). Average lesion area (0.107 mm^2^ RD, 0.092 mm^2^ HFD, 0.132 mm^2^ RD-78c, and 0.117 mm^2^ HFD-78c, as determined by hematoxylin and eosin, Supplemental Figure 1C) was not significantly altered by either HFD (F_(1, 24)_= 2.642, p=0.117) or 78c (F_(1, 24)_=0.886, p=0.356). Neurofilament immunoreactivity (a marker of axon integrity, Supplemental Figure 1D), was 18.09% RD, 15.76% HFD, 18.29% RD-78c, and 18.71% of the lesion area on average, and also remained unchanged by drug (F_(1, 24)_= 0.717, p=0.405) or diet (F_(1, 24)_= 0.261, p=0.614). Overall, lysolecithin mice fed 78c along with either a RD or HFD showed significantly greater numbers of remyelinated axons compared to the lysolecithin mice fed a RD or HFD without 78c (F_(1, 25)_= 10.11, p= 0.004; Figure 4B). In particular, more than double the number of remyelinated axons was present in the 78c-regular diet mice over regular diet alone (p=0.0013). Further supporting a role of CD38 inhibition in myelin repair, MBP immunoreactivity in the total lesion or the lesion rim was higher in the 78c-treated mice irrespective of diet (F_(1, 24)_= 5.71, p=0.025; Figure 4C). Importantly, numbers of Olig2^+^ (F_(1, 23)_ = 9.508, p= 0.005; Figure 4D) and GST3^+^ (F_(1, 24)_= 6.172, p=0.020; Figure 4E) cells were significantly higher in the combined 78c group (which includes LL-RD-78c and LL-HFD-78c mice) versus the combined group without 78c treatment (which includes LL-RD and LL-HFD). Moreover, the number of Olig2^+^ cells in the lesion rim (p=0.049) and GST3^+^ cells in the rim (p= 0.044) or total lesion (p= 0.032) were significantly higher in the LL-RD-78c versus the LL-RD group. These findings suggest that, *in vivo,* 78c helps to preserve or replenish myelinating cells and their progenitors, leading to increased markers of myelin repair. Furthermore, these results provide a strong rationale for targeting CD38 and overall NAD^+^ metabolism as a new strategy to attenuate oligodendrocyte loss and aid in myelin regeneration.

### 78c promotes pro-repair astrocytes after lysolecithin-induced demyelination

Next, we chose to explore the role of 78c on neuroinflammation markers in the spinal cord following demyelination. Histological quantification of threshold-positive staining for microglia marker IBA1 (Figure 5A) indicated a significant interaction between diet and drug treatment on the lesion rim (F_(1, 24)_= 6.936, p=0.015), yet no significant effect of diet (F_(1, 24)_= 2.160, p=0.154) or drug (F_(1, 24)_= 0.2549, p=0.618) alone. Although there is a trend of decrease for both IBA1 and GFAP in the 78c treated group, no statistically significant attenuation of astrocyte marker GFAP was observed (Figure 5 B). In both the RD- and HFD-fed mice, there was, however, a significant increase in S100a10 immunoreactivity with 78c treatment (F_(1, 22)_= 20.77, RD: p=0.001, HFD: p=0.045; Figure 5C), considered a marker for pro-repair astrocytes. There was also a slight decrease in C3d staining, a marker of reactive astrocytes, with 78c treatment, although this did not reach statistical significance (F_(1, 19)_= 3.681, p=0.069; Figure 5D).

**Figure 5.**
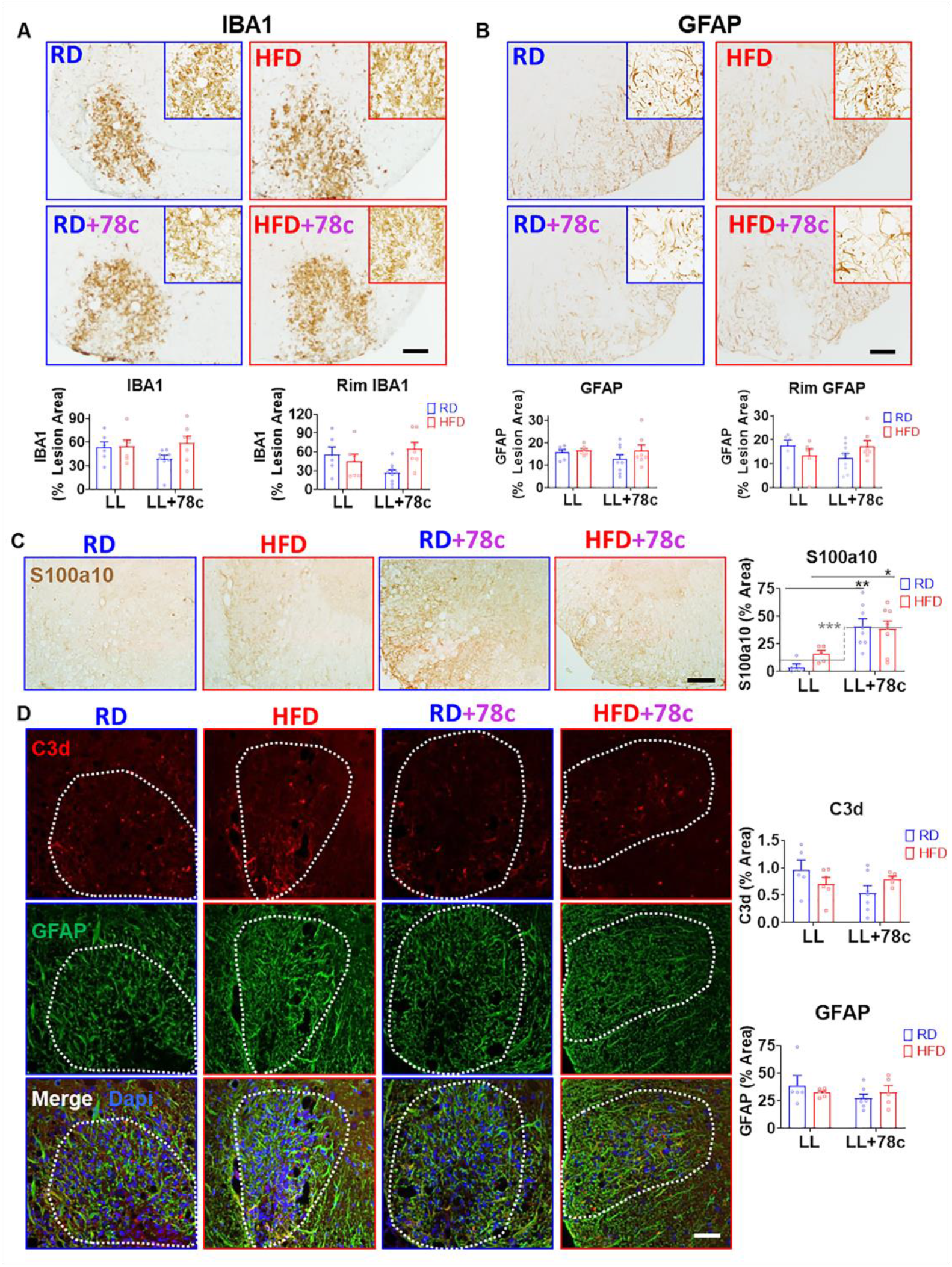
78c promotes pro-repair astrocytes after lysolecithin-induced demyelination. Histological analysis of markers of neuroinflammation in animals fed a regular diet (RD) or high fat diet (HFD) with or without 78c (CD38 inhibitor, 600 ppm) and injected with lysolecithin (LL) followed by a 2-week recovery period. **A, B** IBA1 (**A**, microgliosis) and GFAP (**B**, astrogliosis) were not significantly affected by either diet or 78c alone, although a significant interaction for drug and diet was detected for IBA1 immunoreactivity in the lesion rim. Scale bar= 100 microns. **C-D**, Changes in and pro-repair (S100a10, brown) pro-inflammatory (C3d, red) astrocyte (GFAP, green) markers in adjacent sections within the LL lesion area indicates a shift toward pro-repair in the 78c-treated mice. Scale bar = 100 µm. Bar graphs represent mean +/− SEM. Two-way ANOVA (n=5-8 mice/group). Asterisks indicate levels of statistical significance, where **, p<0.01 when compared to LL-RD by Bonferroni post-test. Gray lines on graphs indicate genotype means, while gray broken lines with asterisks indicate a significant effect of genotype overall.

### Astrocytes cultured in high saturated fat conditions increase CD38 and pro-inflammatory cytokine expression and inhibit oligodendrocyte differentiation

Astrocytes release various growth factors, cytokines, chemokines, extracellular matrix proteins, glutamate, and reactive oxygen species that heavily influence oligodendrocyte proliferation and differentiation (25, 26, 32, 35). Therefore, we next investigated the effects of culturing oligodendrocytes in conditioned media obtained from PA-treated astrocytes with or without 78c co-treatment, as depicted in Figure 6A. As expected, astrocyte-conditioned medium (ACM) from PA-treated astrocytes significantly reduced oligodendrocyte PLP (F_(2, 17)_= 9.396, p=0.006) and MBP expression (F_(3, 8)_= 8.155, p=0.003). However, this negative impact on myelin protein production was partially rescued by co-treatment with the small molecule CD38 inhibitor 78c (3 µM) (p=0.002, PLP; and p=0.040, MBP) (30, 34). Likewise, addition of exogenous NAD^+^ (50 µM) to the PA-treated astrocyte cultures (F_(2, 9)_= 8.502, p=0.008; Figure 6 B) significantly increased PLP expression, while ACM-treated oligodendrocyte numbers remained unchanged across treatment groups (F_(2, 9)_= 1.153, p=0.358). Along with other research that indicates the role of NAD^+^ in reducing astrocyte proinflammatory and oxidative stress responses (18, 36), these data indicate the importance of maintaining proper astrocyte NAD^+^ levels in order to promote oligodendrocyte differentiation.

**Figure 6.**
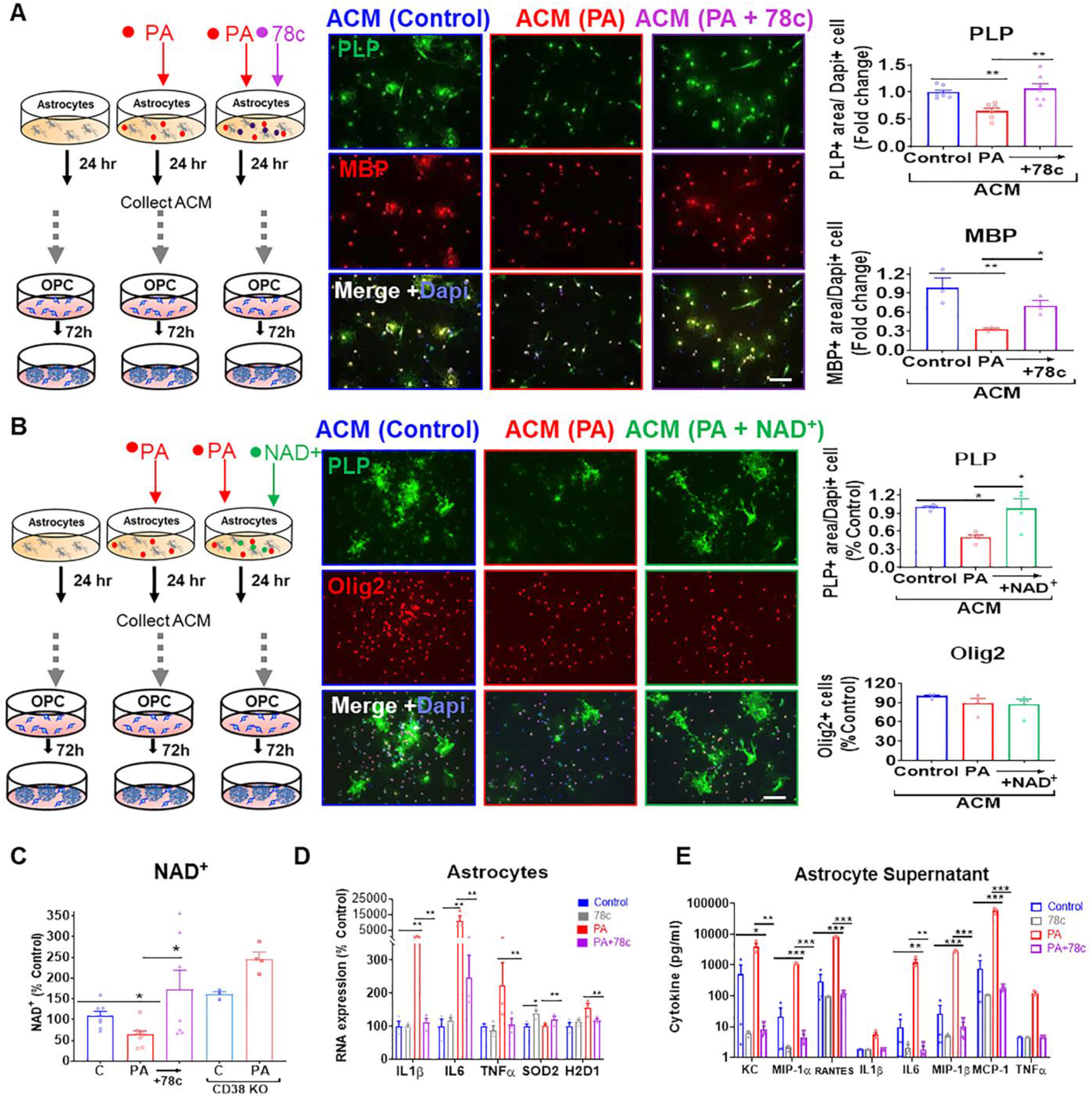
Astrocytes cultured in high saturated fat conditions increase CD38 and pro-inflammatory cytokine expression and inhibit oligodendrocyte differentiation. **A** Schematic depicting how astrocyte conditioned media (ACM) treatments on primary murine oligodendrocytes were performed. Conditioned media obtained from PA-treated murine astrocytes (ACM (PA)) reduced protein expression of mature oligodendrocyte markers PLP and MBP, while PA-78c-ACM-treated OLCs have enhanced PLP and MBP expression. **B** Representational diagram of astrocyte treatments with PA and NAD^+^ to collect ACM for OPC treatments. The addition of NAD^+^ restored PLP levels when compared to ACM-PA, but had no effect on Olig2. Scale bar = 100µm. **C-E** High fat consumption was modeled in primary murine astrocyte cultures by including saturated fat palmitate in the media (PA, 100 µM, 24 h). *C*, Addition of a small molecule CD38 inhibitor (78c, 3µM) was effective to restore NAD^+^ levels in PA-treated astrocytes when assessed using an NAD^+^ cycling assay. Astrocytes derived from CD38 knockout mice did not show depleted NAD^+^ levels in response to PA. RNA levels by qPCR (*D*) of proinflammatory cytokines, antioxidant enzyme, and an astrocyte activation marker in PA and 78c treated astrocytes and cytokine levels (*E)* detected in astrocyte conditioned media (ACM) by Bioplex assay. Bar graphs represent the mean +/− SEM. Asterisks indicate levels of statistical significance when compared to control as determined by or one-way ANOVA with Bonferroni post-test, where *, p<0.05; **, p<0.01; ***, p<0.001 versus control.

To address whether the possible NAD^+^-driven astrocytic changes indirectly mediate the observed effects on oligodendrocyte, we next used primary murine astrocyte cultures to assess NAD^+^ levels and transcriptional changes. Addition of 78c (3µM) was effective to restore NAD^+^ levels in PA-treated astrocytes when quantified using an NAD^+^ cycling assay (F(_4, 23_)= 6.113; p=0.004; Figure 6C). However, astrocytes derived from CD38 knockout mice did not have depleted NAD^+^ levels in response to PA. Next, we determined whether CD38 regulates astrocyte pro-inflammatory responses. After treating astrocyte cultures with PA and/or 78c, we used qRT-PCR and a multiplex bioassay for determining cytokine RNA transcripts and release into culture supernatant, respectively. We observed significant increases in IL-1β and IL-6 expression (F_(3, 8)_= 8.427, p=0.007, IL-1 β; and F_(3, 8)_= 11.83, p=0.007, IL-6; Figure 6D) in astrocytes stimulated with PA and 78c significantly attenuated PA-induced RNA expression of IL-1β (p=0.007), IL-6 (p=0.008), TNFα (p=0.017), and reactive astrocyte marker H2D1 (p=0.028). Additionally, the mitochondrial antioxidant SOD2 was significantly increased in astrocytes treated with 78c (p=0.022) or both PA and 78c (p=0.035). Significant increases in secretion of cytokines triggered by PA were observed using a Bioplex assay including CXCL1 (KC) (F_(3, 8)_ = 9.296, p= 0.014), MIP1α (F_(3, 8)_= 549.8, p<0.0001), RANTES (F_(3, 8)_= 418.5, p<0.0001), IL-6 (F_(3, 8)_= 18.00, p=0.001), MIP1β (F_(3, 8)_= 533.5, p<0.0001), and MCP1 (F_(3, 8)_= 66.96, p<0.0001)(Figure 6 E). Notably, co-treatment with 78c successfully mitigated PA-stimulated secretion of CXCL1 (p=0.006), MIP1α (p<0.0001), RANTES (p<0.0001), IL-6 (p=0.001), MIP1β (p<0.0001), and MCP1 (p<0.0001). Thus, inflammatory and oxidative stress-related changes in cultures astrocytes may help to explain reduced differentiation potential of the oligodendrocyte progenitors in the presence of PA-treated astrocyte conditioned medium.

### NAD**^+^** and NMN restore oligodendrocyte differentiation in PA-treated cultures

Our recent findings demonstrated that PA can directly impair oligodendrocyte differentiation from neural stem cells or oligodendrocyte progenitors (4), and others have indicated that depletions in cellular NAD^+^ can prevent differentiation of stem cells to mature oligodendrocytes (37). To examine the effects of replenishing NAD^+^ levels on oligodendrocytes, we next supplemented cultures with exogenous NAD^+^ (50 µM) and monitored PLP expression (Figure 7A). Co-treatment with PA and NAD^+^ significantly enhanced PLP expression (F_(2, 16)_=3.458, p=0.024; Figure 7B) compared to exclusive PA treatment. However, treatment of oligodendrocytes with CD38 inhibitor 78c did not attenuate this effect (data not shown).

**Figure 7.**
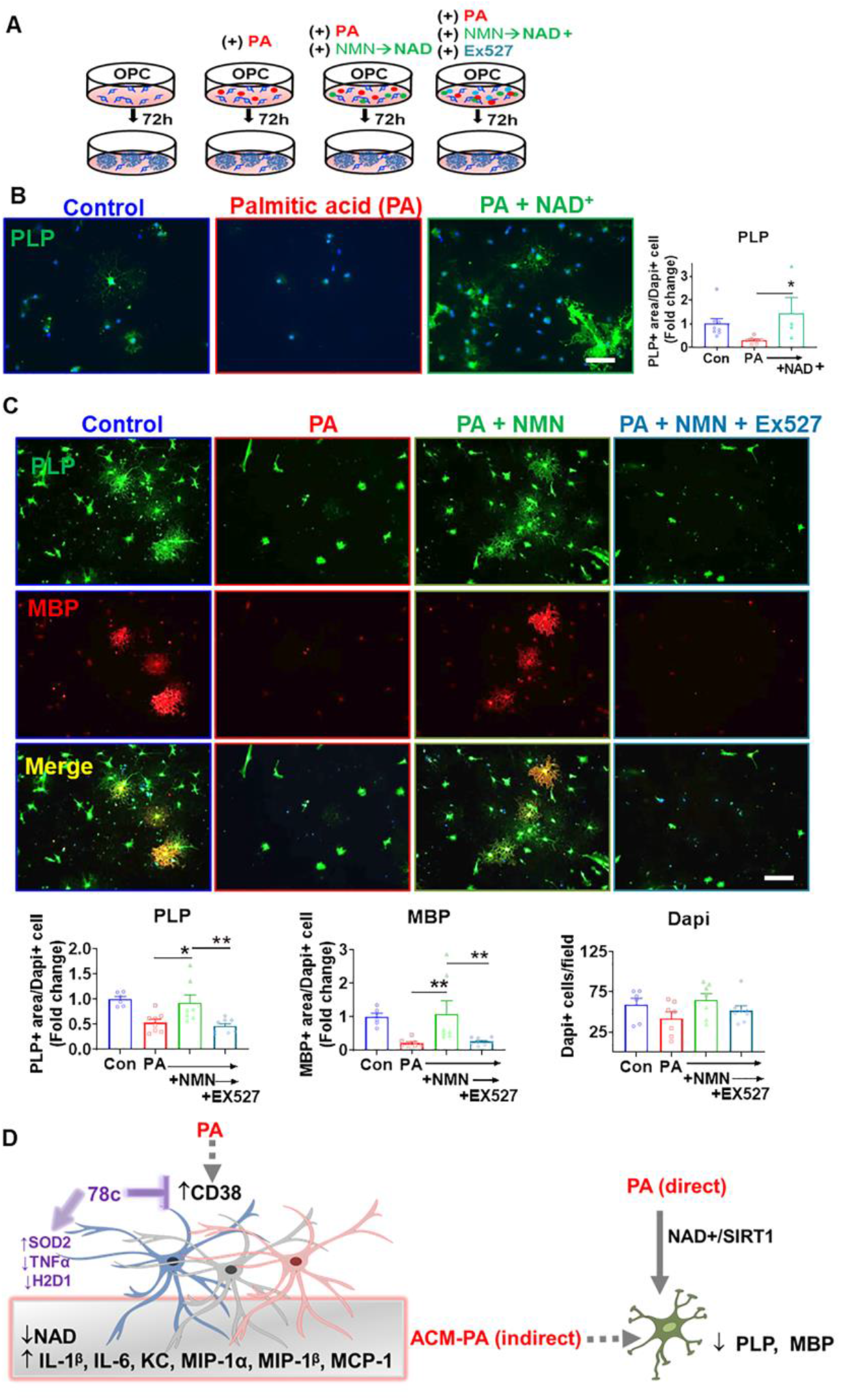
NAD^+^ replenishment attenuates PA-induced deficits in oligodendrocyte differentiation in a SIRT1-dependent manner. **A** Representational diagram of experimental treatments in primary murine oligodendrocyte lineage cells (OLCs). Adding exogenous NAD^+^ (50 µM) to oligodendrocyte cultures significantly restores PLP protein expression (***B***) in PA-exposed OLCs. **C** Treatment with NAD^+^ precursor nicotinamide mononucleotide (NMN,100 µM) also enhances expression of mature oligodendrocyte markers in PA-treated cultures, although not when SIRT1 inhibitor Ex-527 (5 µM) is also included in the media, as determined by immunostaining quantification for PLP and MBP per DAPI^+^ cell. Scale bar= 100 µm. **D** Schematic summarizing direct and indirect effects of palmitate on oligodendrocytes characterized *in vitro*. Graphical results represented as the mean ± SEM (n=4-8/group). Asterisks indicate levels of statistical significance when compared to designated group as determined by one-way ANOVA with Bonferroni post-test, where *, p<0.05; **, p<0.01; ***, p<0.001.

Interestingly, CD38^+^ cells can deplete NAD^+^ and metabolites available for low-CD38 expressing cells (30, 38), such as oligodendrocytes (16–18, 28). We found that direct effects of saturated fat treatment on oligodendrocytes include mitochondria function and morphology changes (4) while other mechanisms, potentially including NAD^+^-depleting mechanisms (37, 39, 40), could also play a part. SIRT1 regulates mitochondria function, was decreased following HFD in spinal cord tissue (4), and is a NAD^+^-dependent deacetylase. We hypothesized that the pro-myelinating effects of increasing NAD^+^ are SIRT1-dependent, so we determined if the beneficial effects of NAD^+^ in the presence of PA could be blocked with a SIRT1 inhibitor Ex-527. Similar to adding exogenous NAD^+^, treatment with the NAD^+^ precursor NMN (100 µM) also restored markers of oligodendrocyte differentiation in PA-treated cultures (F_(3, 25)_=8.449, p=0.011; Figure 7C). However, in the presence of Ex-527 (5 µM), the increases in PLP and MBP protein in response to NMN were suppressed (p=0.003). At the doses used, PA, NMN, and Ex-527 did not significantly alter the number of oligodendrocytes. Hence, the impaired ability of oligodendrocytes cultured with excess saturated fat to differentiate is at least partially regulated by NAD^+^-dependent sirtuin activity (Figure 7D).

### 78c prevents lysolecithin- and PA-mediated myelin protein and oligodendrocyte loss

Since we observed protective effects of genetic CD38 inhibition on oligodendrocyte survival following HFD, as well as increased CD38 expression in a spinal cord demyelination model, we next sought to explore benefits conferred by pharmacologic inhibition using 78c in a brain model of lysolecithin-induced *ex vivo* demyelination. Murine cerebellar slices from postnatal pups were demyelinated using lysolecithin (LL, 0.5 mg/ml, 18 h), followed by a period of one week to allow for repair (Figure 8 A). Confocal imaging of immunofluorescence staining demonstrated reductions in MBP after LL that was significantly attenuated by 78c co-treatment (F_(2, 25)_=4.578, p=0.020; Figure 8B).

**Figure 8.**
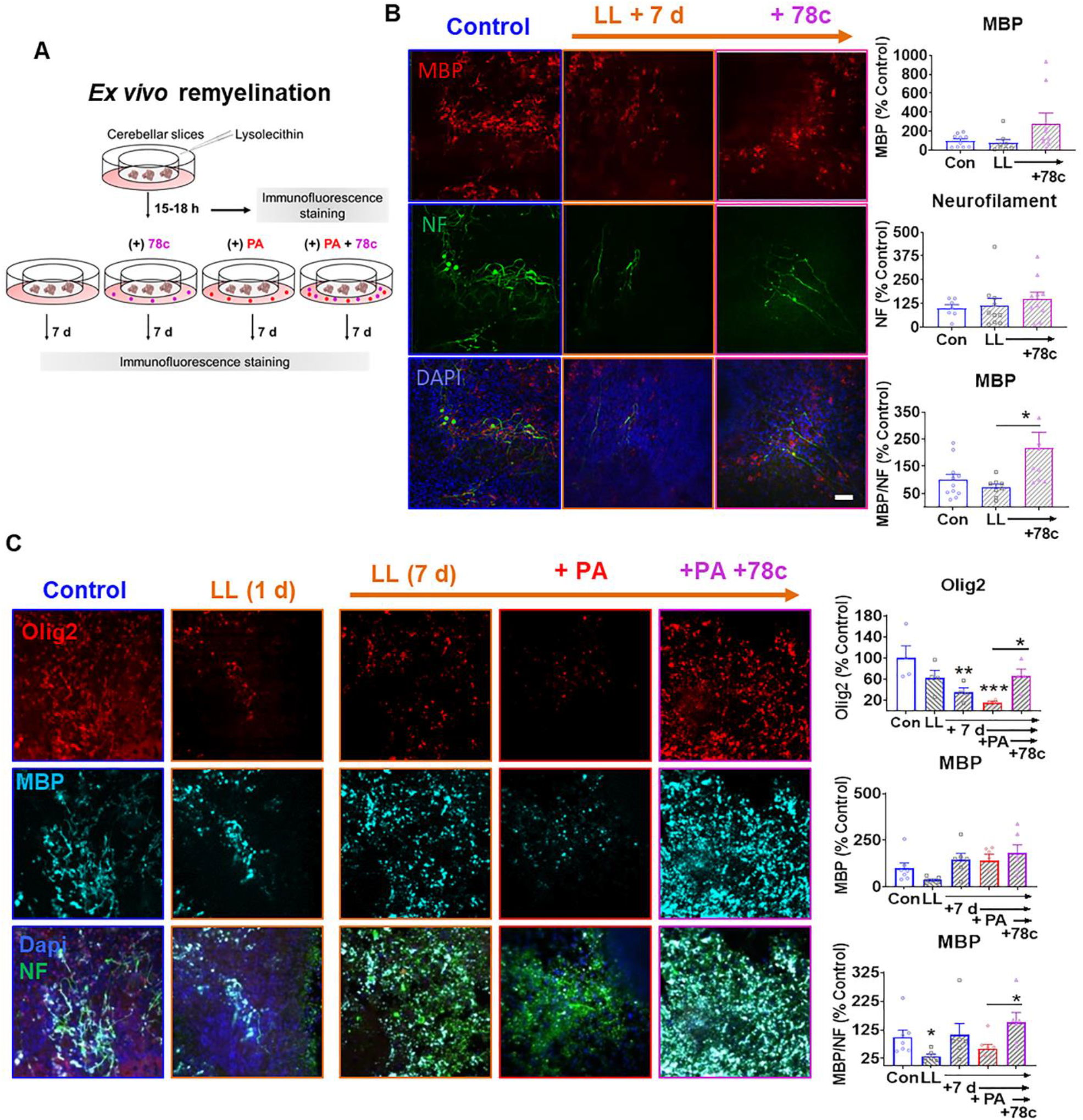
CD38 inhibition protects from saturated fat and lysolecithin-induced demyelination in cerebellar slices. **A-B** Representational diagram (**A**) of *ex vivo* remyelination experiments where murine organotypic cerebellar slices were demyelinated with lysolecithin (LL, 0.5 mg/ml, overnight) followed by subsequent culture in control media for 1 wk. Confocal microscopic imaging of slices indicate reduced MBP (red, ***B***) expression when normalized to neurofilament (NF, green). LL-demyelinated slices cultured with 78c had significantly increased MBP expression when normalized to NF staining and expressed as a percentage of the control group. **C** Slices cultured in media containing both LL-palmitic acid (PA, 100 µM) and 78c, a CD38 inhibitor, 3 µM) had significant improvements in Olig2 and MBP expression compared to the LL-PA alone group. Bar graphs represent the mean ± SEM (n=4-8/group); *, p<0.05, **, p<0.01, and ***, p<0.001 when compared to control by one-way ANOVA with Bonferroni post-test.

Finally, to evaluate the therapeutic potential of 78c in a brain model of high fat exposure combined with demyelination, organotypic cerebellar slices were demyelinated as described above and cultured for 7 d in the presence or absence of PA and/or 78c. Cerebellar slices showed fewer Olig2^+^ cells (F_(6, 21)_=4.964, p=0.003) with LL insult. Inclusion of PA in the media further diminished Olig2+ cells in the LL slices. After 7 d of recovery, Olig2^+^ cells remained significantly depleted in the 7 d LL group. There was also less MBP immunoreactivity following LL (F_(6, 21)_=2.899, p=0.020; Figure 8 C), however MBP levels were successfully restored after 7 d recovery. Importantly, 78c and PA co-treatment significantly increased the percentage of Olig2^+^ cells (p=0.049) and MBP expression (p=0.023) in the demyelinated cerebellar slices. However, neither palmitate nor 78c alone, without lysolecithin, for two weeks significantly altered MBP expression (PA, p=0.668, and 78c, p=0.566 versus control). These findings demonstrate the capability of 78c to restore myelin protein expression even in a combined saturated fat- and demyelinating agent-induced condition. Overall, these data suggest that 78c modulates astrocyte responses elicited by demyelinating injury or high fat diet in the C NS, skewing them towards a phenotype that enhances oligodendrocyte survival and differentiation and myelin repair (Figure 9).

**Figure 9.**
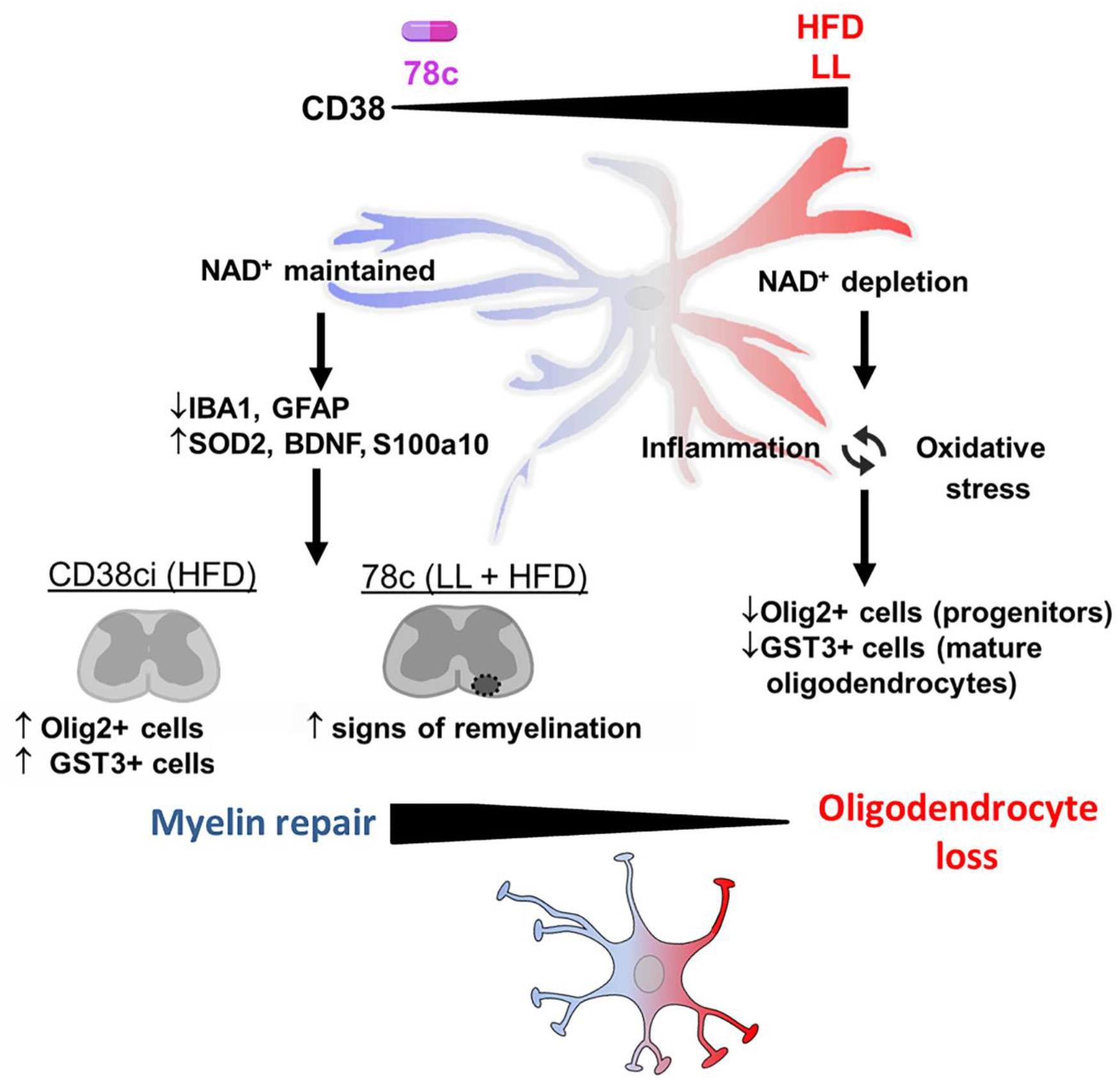
Schematic representation of proposed hypothesis. Resting astrocytes typically help to provide a supportive environment for oligodendrocyte differentiation and repair, when necessary. High fat diet (HFD) and lysolecithin (LL) increase CD38 expression in reactive astrocytes, thereby depleting NAD^+^ levels. Reductions in NAD^+^ result in oxidative stress and inflammation which triggers impaired differentiation and survival in oligodendrocyte lineage cells. 78c inhibits CD38 to attenuate the described detrimental effects of astrocytes on oligodendrocytes.

## Discussion

### Relevance of Diet and NAD^+^ homeostasis to Multiple Sclerosis

Myelin disturbances and oligodendrocyte loss can leave axons vulnerable leading to permanent neurological deficits. The results of the current study suggest that metabolic disturbances, triggered by consumption of a diet high in fat, promote oligodendrogliopathy and impair myelin regeneration through astrocyte-linked indirect NAD^+^-dependent mechanisms. Diet-induced obesity increases the risk for developing MS, especially when combined with other known risk factors (1, 2). We demonstrate that restoring NAD^+^ levels via genetic inactivation of CD38 can overcome these effects. Moreover, we show that therapeutic inactivation of CD38 can enhance myelin regeneration. Together these findings point to a new metabolic targeting strategy positioned to improve disease course in MS and other conditions in which the integrity of myelin is a key concern.

### Neuroinflammation and CD38

Neuroinflammation and astrogliosis are predominant pathological mediators of neurodegenerative conditions, including MS (9, 41), and astrogliosis is linked to HFD-induced CNS pathology in rodents (8, 24). In spinal cord astrocytes of HFD mice, we observed increased CD38 expression and an overall depletion of NAD^+^. These findings are consistent with our human immunostaining data and other reports of increased CD38 in the brain of an individual with MS (42) and in experimental models of neurological injury and disease (21). Known upstream activators of CD38 expression include cytokines, lipopolysaccharide (LPS), 25-hydroxycholesterol (25-HC), glutamate, and factors secreted by senescent cells (38, 43–45). In the current study, we found that HFD or focal lysolecithin-mediated demyelinating injury each drove increased CD38 expression in astrocytes, while genetic or pharmacological CD38 inhibition after HFD or demyelination, respectively, attenuated astrocyte activation markers. Confirming the human relevance of this effect, we observed CD38 in hypertrophic reactive astrocytes within an active demyelinating lesion from an individual with MS.

Astrocytes and microglia secrete a variety of chemokines, cytokines and growth factors that regulate myelin injury and regeneration (25, 35). In palmitate-stimulated astrocytes, IL-6 and other cytokines were increased. CCL3/MIP1α, MIP1β, and RANTES also increase in active demyelinating MS lesions and fluids (46, 47). Pointing to a therapeutic strategy, this response was blunted when NAD^+^ levels were restored by co-treatment with 78c. Other studies indicate that CD38 inhibition or knockout decreases cytokine secretion in LPS-stimulated microglia, macrophages, and monocytes (48). Recently, knockout of CD38 or supplementation with NAD^+^ was shown to suppress activation of astrocytes in culture (18). Therefore, improving NAD^+^ levels by targeting CD38 shows therapeutic promise across multiple neurological and non-neurological disease types (21), and here we establish its significance to oligodendrocyte heath and regeneration with strong linkage to its effects on astrocyte metabolism.

### Oxidative stress and oligodendrocyte differentiation

Reactive oxygen species can also inhibit oligodendrocyte survival (32). Complementing this, we demonstrate more SOD2 and less IL1β in high fat-78c co-treated astrocyte cultures compared to treatment with high fat alone. We also found more SOD2, BDNF and S100a10 in the CD38ci mouse spinal cord pointing to a shift towards pro-repair astrocyte properties (9). Blocking downstream CD38 signaling also reduced markers of oxidative damage in a convulsion model (49). Our findings suggest that the beneficial effects of CD38 inhibition include protection against astrocyte pro-inflammatory factors and reactive oxygen species known to negatively impact oligodendrocyte renewal and maturation.

The effects of CD38 activity and NAD^+^ levels may depend heavily on cell type and disease context, with findings here supporting a critical protective role of lowering astrocyte CD38 activity for protection of myelinating cells and for myelin regeneration in the adult CNS. Future studies could further solidify this finding once astrocyte-specific conditional knockouts for CD38 become available. CD38-positive cells can regulate NAD^+^ levels in neighboring CD38-negative cells by influencing NAD^+^ precursor availability (30, 38), which may be the case for astrocytes and oligodendrocytes in our studies. Consistent with this model, 78c had no direct effect on oligodendrocyte differentiation in the context of palmitate *in vitro*. Palmitate can inhibit nicotinamide phosphoribosyltransperase (NAMPT) activity (40), a rate-limiting enzyme in the NAD^+^ salvage pathway, as well which prevents differentiation of stem cells to oligodendrocyte lineage cells (37) and potentially explains the direct inhibitory effect of palmitate on oligodendrocyte differentiation observed here. Thus, complex alterations in NAD^+^ biosynthesis, salvage, and degradation pathways may contribute to the effects of palmitate on oligodendrocytes that we document can be rescued by NAD^+^ replenishment.

### NAD^+^-boosting strategies for neural repair

Several recent studies have explored supplementing NAD^+^ precursors or flavonoids known to inhibit NAD^+^ase activity to improve myelin repair, neuroprotection, and to decrease immune cellular responses. For example, an NAD^+^/BDNF/TrkB pathway is proposed to be involved in nicotinamide-mediated protection, including improved remyelination in a stroke model (50) and precursor supplementation increased BDNF in a Huntington’s disease model (51). Supporting a possible NAD^+^-BDNF pathway for remyelination, we observed increased BDNF expression in the spinal cord of CD38ci mice and improved remyelination with oral 78c administration. Although direct effects on myelinating cells were not examined, mice with CD38-knockout, given the CD38 inhibitor apigenin, or supplemented with NAD^+^ each showed reduced EAE disease severity (15, 52, 53). NAD^+^ levels can impact oligodendrocyte lineage cell proliferation and survival via cyclins and E2f1, while differentiation is reported to be regulated redundantly by SIRT1 and SIRT2 (37, 54, 55). Results here demonstrate that the ability of NAD^+^ supplementation to promote oligodendrocyte differentiation *in vitro* is indeed at least partially reliant on SIRT1 function. Restoring NAD^+^ levels was also beneficial in the context of palmitate or high fat-induced inflammation and oxidative stress via SIRT1 in liver, heart, and muscle (56–58). Improvements in expression of myelin proteins in the spinal cord of mice consuming a high fat high sucrose diet in conjunction with coordinate wheel running, were also linked to increases in SIRT1 expression (6). Collectively, these findings are particularly encouraging since effective modulation of disease progression will likely include a combination of therapeutic strategies encompassing anti-inflammatory, myelin regeneration, and neuroprotective interventions.

### Study considerations

HFD consumption did not further impair markers of myelin integrity in our *in vivo* model of myelin injury and repair as it did in *ex vivo* remyelinating cerebellar slice cultures. Although treatment with 78c did improve myelin regeneration in mice consuming a regular diet, little impact was seen in mice consuming a HFD, suggesting the need to explore additional factors in the context of HFD-demyelination that contribute to impairments. Alternately, a higher dose or combination of drugs (to target various aspects of astrocyte responses) may be required to improve remyelination in the context of HFD. Although a preference for pro-repair versus pro-inflammatory phenotype was indicated following 78c treatment in the lysolecithin model of focal demyelination studied, a complete astrocyte-specific transcriptomic profile in the HFD mice with demyelination will be needed to identify novel genes in the astrocyte transcriptome that could be targeted to mitigate the indirect oligotoxic effects we observe.

### Conclusions

Our data demonstrates an essential role of NAD^+^ levels in maintaining CNS white matter integrity, which can be impaired by focal demyelination or high fat diet consumption. Further, we have shown that inhibition of the NADase CD38 attenuates oligodendrocyte loss and promotes myelin repair. Genetic CD38 inhibition attenuated HFD-induced oligodendrocyte loss and dietary supplementation with 78c increased counts of oligodendrocytes and remyelinated axons following focal demyelinating injury. These findings suggest that increases in astrocyte CD38, and the associated depletion of NAD^+^ levels, may contribute indirectly to impairments in oligodendrocyte health and myelin regeneration via inflammation and oxidative stress. Tarrago, et al. (30) recently established 78c as a specific, reversible, and non-competitive inhibitor of CD38 NAD^+^ase activity improves age-related metabolic function (28). Altogether, these findings suggest that genetic or pharmacologic CD38 inhibition provides therapeutic benefits in HFD-induced metabolic dysfunction and downstream signaling events in the CNS, in addition to the already described peripheral benefits on age-related metabolic dysfunction. Importantly, inhibition of CD38 with 78c can improve myelin regeneration in the adult CNS including reducing the pro-inflammatory profile while increasing the pro-repair profile of astrocytes. Together these findings point to the potential translational value of targeting CD38 to improve outcomes in MS and other demyelinating conditions.

A few current MS disease modifying therapies (i.e. glatiramer acetate, fingolimod, dimethyl fumarate) affect CD38^+^ cell counts or inhibit astrocyte activation, but these roles have not been directly linked to therapeutic benefits (11, 15, 59–62). For instance, fingolimod treatment decreased CD38 expression in a rat EAE model (15), and was shown to promote beneficial astrocyte responses in culture (63). Monoclonal antibodies targeting CD38 have already met with success in clinical trials for conditions outside the CNS (43, 64–66), including Daratumumab which is FDA approved for use in multiple myeloma. Accordingly, the research findings we present should inspire further research into ways to mitigate the negative consequences of HFD on oligodendrocytes alone and in the context of white matter injury which may include dietary intake of NAD^+^ precursors, administration of CD38 inhibitors, or exercise-related rehabilitation strategies.

## Materials and Methods

See Table 1 for key reagents.

**Table 1.**
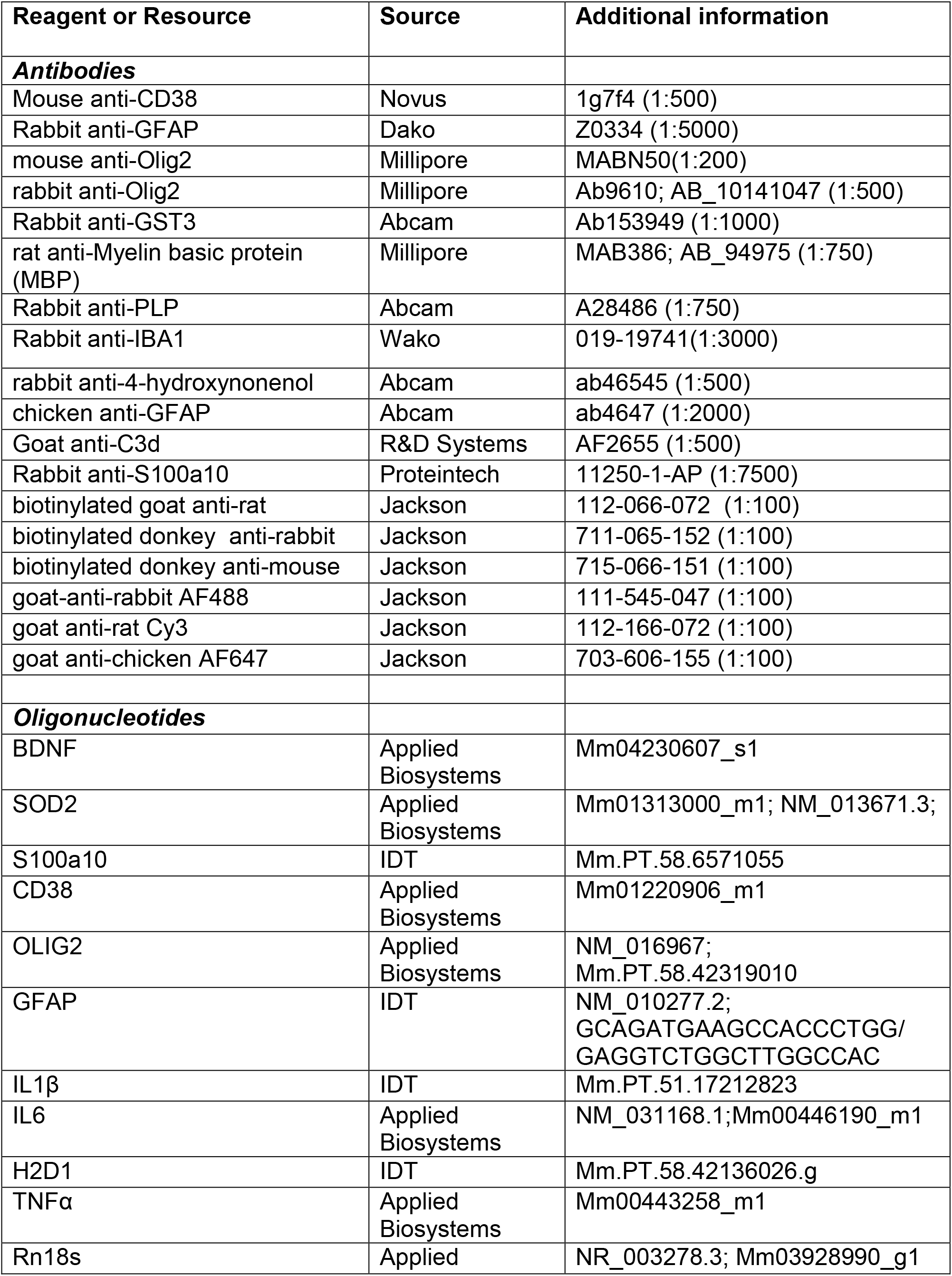
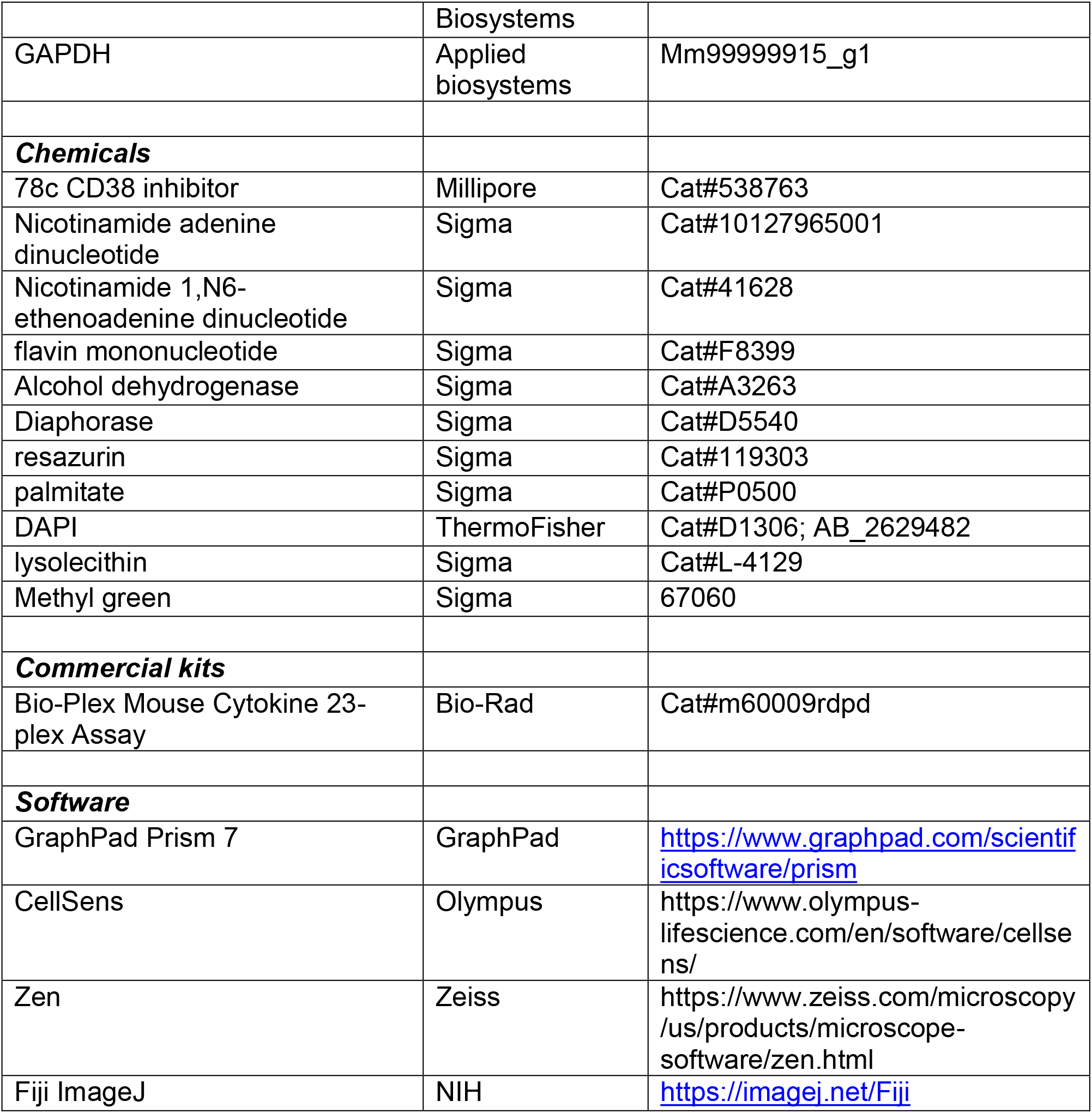
Key Resources Table

## Animal husbandry and diets

All experimental diets and procedures were approved by The Mayo Clinic Institutional Animal Care and Use Committee (IACUC) and performed in accordance with institutional and regulatory guidelines. Mice were provided *ad libitum* access to diets along with tap water. For initial characterization of CD38 expression and NAD^+^ levels following a high-fat diet, male C57BL6/J mice were obtained from Jackson Labs (Bar Harbor, ME) and were randomly assigned to feeding on either a regular diet (RD, 10 % kcal fat, D12450K, Research Diets, New Brunswick, NJ) or a high-fat diet (HFD, 60% kcal fat, D12492, Research Diets, New Brunswick, NJ) beginning at 10 wk of age, as previously described (4).

For studies comparing wildtype and CD38ci mice fed RD and HFDs, mice were bred onsite after initial generation at TransViragen (Chapel Hill, NC) via recombineering technology, as previously described (30, 67). Male mice received a regular (10.4% kcal fat, Teklad TD000357) or HFD (42% kcal fat, Teklad TD88137) beginning at 50 wk of age for 30 wk. Diets used for lysolecithin demyelination studies were these same diets, with or without 600 ppm 78c added (Calico Life Sciences, LLC), and were started one week prior to the spinal cord microinjection procedure. Food intake and body weights were monitored weekly.

### Lysolecithin microinjection into spinal cord

For focal demyelination of the ventral white matter (laminectomy at thoracic 10^th^ vertebra; mediolateral 0.3, dorsoventral −1.4 mm relative to the dorsal surface) of the murine spinal cord, 12 wk old male mice were injected with 2 µl of a saline solution containing 1% lysophosphatidyl choline with 0.01% Evan’s Blue (Sigma L4129) at 0.2 µl/min using a 30-40 µm glass micropipette and stereotaxic microinjection device (Stoelting, Inc., Wood Dale, IL). Male mice were selected for lysolecithin injection procedures to reduce variability due to progesterone-mediated enhancement of myelin repair and modulation of CNS neuroinflammatory responses (68). Young adult mice have previously yielded reliable, reproducible lesions and good survival in this model from studies in our laboratory and others (23, 69). The spinal cord has large white matter tracts that are more frequently myelinated axons versus corpus callosum, allowing for more confident quantification of remyelinated axons due to challenges of determining thin myelin sheaths or simply unmyelinated axons from demyelinated axons (23, 69, 70).

For anesthesia, 100 mg/kg ketamine and 10 mg/kg xylazine were administered intraperitoneally prior to surgery, while postoperative care consisted of an antibiotic (Baytril 50 mg/kg/day, i.p.), an analgesic (Buprinex 0.05 mg/kg, twice daily, subcutaneous) and saline (0.2ml/d, intraperitoneal) for three days. After a two-week recovery period, mice were deeply anesthetized (ketamine 150mg/kg and xylazine 15 mg/kg, i.p.) and transcardially perfused with 4% paraformaldehyde (Fisher T353-500) for collection of spinal cord tissue. Spinal cord segments were microdissected, with the rostral 2 mm segment processed for paraffin-embedded blocks and the caudal segment osmicated (1% osmium tetroxide, 0223B, Polysciences) and embedded in Araldite. To identify myelin sheaths, Araldite blocks were cut to 1 µm sections, stained with p-phenylenediamine (Sigma P6001), imaged with 60x objective (Olympus BX51 microscope), and stitched together (Adobe Photoshop) to comprise the entire lesion within the stained tissue section (23, 70). Number of remyelinated axons per square millimeter of lesion was reported after blinded counting in ImageJ (23, 70). Immunohistochemical analysis of focal demyelinated lesions utilized overall neuropathological changes (hematoxylin and eosin) along with myelin loss (based on MBP staining) to determine the lesion core, while the lesion rim was defined as the region encompassing the entire border of neuroinflammatory markers (GFAP, IBA1) (70). Lesion areas between 50,000 - 180,000 µm2 were included in the analysis based on our previous studies using this model (23, 70). Analyses were performed with experimenter blinded to treatment group by assigning a random animal number.

### Immunohistochemistry

Spinal cords were fixed in 4% paraformaldehyde, processed, paraffin-embedded, and cut to 6 µm for all immunoperoxidase and immunofluorescence procedures, as described previously (4, 23). When necessary (all antibodies except GFAP and MBP), heat-induced antigen retrieval was achieved through incubation with sodium citrate (10 mM, pH 6.0) in a steamer basket. Sections were blocked (20 % v/v normal goat serum in PBS), then incubated in primary antibodies described in the key resources table.

The following day, incubation with secondary antibodies (biotinylated or fluorochrome-conjugated, Jackson, 1:200) for immunoperoxidase or immunofluorescence staining was performed. After incubating in an avidin peroxidase solution (Vectastain Elite ABC kit, PK-6100, Vector Laboratories), color development was achieved by incubation with 3,3-diaminobenzidine (DAB, Sigma D5637) and counterstaining with methyl green (Sigma, 67060). Slides were dehydrated, cover slipped with DPX (Sigma), and images acquired using and Olympus BX51 microscope with DP72 camera. Immunofluorescence slides were counterstained with 4’,6’- diamindino-2-phenylindole (DAPI), mounted with fluoromount, and imaged with an Olympus BX51 microscope and captured with Olympus XM10 camera. Cell and axon counting, as well as thresholding of images, was performed in ImageJ (NIH) without knowledge of treatment groups using cropped regions of interest (Allen Spinal Cord Atlas, Adobe Photoshop) and averaged from two sections per animal (60 µm apart).

Formalin-fixed paraffin-embedded 5 µm thick sections from confirmed MS and normal CNS control autopsy cases were immunostained for CD38 (1:2000, LifeSpan BioSciences, LS-C817841). Steamed antigen retrieval was performed with DAKO high pH target retrieval solution (K8004, Dako) (71).

### RNA isolation and qRT-PCR

To determine gene expression from spinal cord or primary murine cells, total RNA was extracted using RNA Stat60 (Tel-Test; Friendswood, TX) per manufacturer’s instructions and used for qRT-PCR using BioRad iTaq Universal Probes or primers and SYBR Green One-step kits. Starting quantity for quantification of each gene’s expression was normalized to Rn18s or GAPDH. Detailed primer and probe information are annotated in the key resources Table.

### Mixed glial preparations and primary cell cultures

Primary mixed glial cultures were prepared from P0-3 mice, as previously described (4, 23). In brief, dissected cerebral cortices were dissociated and grown in DMEM (Fisher 11960-044) with 1 mM sodium pyruvate (Corning 11360070), 20 mM HEPES (Gibco 15630-080), 100 U/ml PenStrep (Life Technologies 15140122), 10% fetal bovine serum (FBS, Atlanta 337 Biologicals S11150), and 5 µg/ml insulin. Following 10-12 days in culture, sequential immunopanning was performed to separate astrocyte, microglia, and oligodendrocyte progenitor cells by agitating at 225 rpm on an orbital shaker (Innova 2000). Purity of the cell types cultured was within the ranges reported previously by our group (72, 73). After removing microglia via differential adhesion, purified oligodendrocyte progenitors were plated onto PLL-coated glass coverslips and cultured in differentiation medium (Neurobasal A media (Life Technologies, 10888022) with N2 (Gibco 17502048), B27 (Gibco 17504044), PenStrep, sodium pyruvate, 2 mM glutamax (Gibco 35050-061), 40 ng/ml T3, and 5% BSA (Sigma A4503)) for two days. Treatments with palmitate (PA, 100 µM) NAD^+^ (50 µM), NMN (100 µM), EX527 (5 µM) and/or 78c (3 µM) were added and cultures were allowed to differentiate for 72 hr. Doses used in cell culture experiments were based on previous studies (30, 74) and preliminary dose-response experiments (data not shown). Positive staining for myelin proteins was compared to control levels by immunofluorescence staining and threshold quantification in ImageJ. Briefly, a positive threshold for each stain was decided, then applied to each image (5 per coverslip) to determine percentage of field positive. Mean values were graphed as a percentage of the control group.

In experiments characterizing astrocyte responses and effects of astrocyte-conditioned media on oligodendrocytes, astrocytes were plated at a density of 450,000 cells/well on PLL-coated 6-well plates and grown for 2-3 days in serum-free Neurobasal A media containing 1% N2, 2% B27, P/S, 1mM sodium pyruvate, 0.45% glucose, 5% BSA, and 50µM β-mercaptoethanol before treating with or without palmitate (PA, 100 µM) and 78c (3 µM). Astrocytes were collected for an NAD^+^ cycling assay or RNA isolation. Supernatants from treated astrocytes were collected 24 h post-treatment and concentrated using Vivaspin sample concentrators (10 and 100 kDa molecular weight cutoffs, Sigma) and stored frozen as 10x protein concentrate until use for treating OLCs (9, 23). Conditioned medium from astrocytes (ACM) was brought back to 1x concentration in oligodendrocyte differentiation medium and placed on oligodendrocytes to differentiate as described above.

### NAD**^+^** cycling and CD38 activity assays

To determine intracellular NAD^+^ levels, one well of a 6-well plate of astrocytes, or approximately 20 mg of spinal cord tissue, was homogenized in 300 µl of 10% trichloroacetic acid in water. Samples were extracted using an organic solvent mixture (1,1,2-Trichloro-1,2,2-trifluoroethane and triocylamine), diluted, and measured along with a standard curve, as previously described (30). Briefly, a cycling reagent (0.76% ethanol, 4 µM FMN, 27.2 U/ml alcohol dehydrogenase, 0.44 U/ml diaphorase, and 8 µM resazurin (Sigma)) was prepared in phosphate buffer (100 mM NaH_2_PO_4_/Na_2_HPO_4_, pH 8.0) and added to an equal volume of pre-diluted cell or tissue extract. Plates were measured every 30 seconds for 1 h at 544 nm using a SpectraMax plate reader. Raw values were compared to a standard curve, then divided by protein quantification (Bradford reagent, Biorad) to be expressed as nmols/mg protein. For measuring CD38 NAD^+^ase activity, spinal cord tissue was homogenized in NETN buffer (100 mM NaCl, 1 mM EDTA, 20 mM Tris-HCl, and 0.5% Nonidet P-40 (Sigma)) supplemented with 5 mM NaF (Sigma) and protease and phosphatase inhibitors (Thermo Fisher) (30). After centrifugation (12000 rpm, 10 min), supernatant volumes were adjusted to protein concentrations and reaction mix containing 50 µM 1,N6-Etheno NAD^+^ in 0.25 M sucrose buffer was added at an equal volume just prior to reading the plate at 300 nm excitation and 410 nm emission every minute for 1 hr on a Spark Magellan plate reader.

### Bioplex assay for cytokines

Astrocyte supernatants from cell treatments described above were used for determination of secreted cytokine levels using a Bio-Plex Mouse cytokine 23-plex assay (BioRad m60009rdpd) per manufacturer instructions. Briefly, post-treatment medium was collected and incubated with primary antibody-conjugated magnetic beads for 30 min. Wells were washed thrice and then incubated for 30 min with detection antibodies. After washing, samples were incubated for 10 min with a streptavidin-phycoerythrin solution, and then washed again prior to resuspension and reading fluorescence with a BioPlex 200 (BioRad) reader and Bio-Plex Manager software.

### Organotypic cerebellar slice culture and demyelination

Organotypic cerebellar slice cultures were cultured and demyelinated using previously described methods (8). Cerebella were dissected from postnatal day 7-9 pups, embedded in low-melting point agar (Invitrogen 16520050), and 350 µm slices collected into artificial CSF solution using a McIlwain tissue chopper (Mickle Laboratory Engineering Co; Guildford, UK). Four slices were transferred and grown per each Millicell insert (Sigma PIHP03050) in 50% MEM, 22.5% Hank’s Balanced Salt Solution (HBSS), 24% heat-inactivated (HI) Horse Serum, 1.5% Glucose, 1% GlutaMax, 1% P/S medium.

Demyelination was induced after 7 d in culture by treating with 0.5 mg/ml lysophosphatidycholine (LL, Sigma) for 18 h. Organotypic slices were placed back in LL-free medium and collected for immunohistochemistry and confocal imaging after 24 h (demyelination) or 7 d (myelin regeneration) with or without addition of PA (100 µM) and/or 78c (3 µM) added to the growth media. Confocal images were acquired at 20x magnification from 5 regions randomly selected per slice using a Zeiss LSM 780 and Zen Software in the Mayo Clinic Microscopy and Cell Analysis Core. Images were thresholded for each stain, and stain-positive immunoreactivity area or cell counts per total field area were quantified in ImageJ.

### Statistics

Data were graphed with bars representing mean ± SEM with sample size for each experiment indicated in the figure legends. To determine sample size, power analysis using the fpower macro in SAS was used to determine animal numbers for each endpoint based on 80% power, α=0.05, and 4 treatment groups. Differences and SD used to determine δ were used from analysis of previous similarly-designed studies in our laboratory. Cell culture studies used a combination of male and female pups, were repeated twice, and assays were run in technical duplicate or triplicate. No statistical outliers were removed, but the ROUT test in GraphPad Prism was used to check for outliers and data normality was assessed using the Shapiro-Wilk test. Comparisons between two groups were analyzed by Student’s t-test, whereas one-way or two-way ANOVAs with Bonferroni post-test were applied for multiple-comparisons. Analysis was performed in GraphPad Prism 7.0 software with p-values ≤ 0.05 considered significant.

### Study Approval

All animal experiments were approved by the IACUC and the human pathology study was approved by the Institutional Review Board of Mayo Clinic, Rochester, MN (IRB 2067-99).

## Author Contributions

IAS and MRL conceived and designed the study. MRL, CC, and TRPM conducted *in vivo* experiments and necropsies. Embedding and histological staining of mouse tissues was performed by MRL and LK. Image acquisition and quantifications were performed by MRL under guidance from CC. YG and CL performed and analyzed human histological staining. MRL completed *in vitro* assays and qRT-PCR. WS and HY assisted with glial preparations and organotypic slice cultures. TRP, CCSC, and ENC provided expertise in NAD^+^ quantification and experimental design as well as CD38 mice. MRL and IAS wrote the manuscript and all authors reviewed and edited the manuscript.

## Acknowledgements

The research was supported by the Mayo Clinic Center for Multiple Sclerosis and Autoimmune Neurology, the Eugene and Marcia Applebaum Foundation, and the Mayo Clinic Center for Biomedical Discovery. Portions of this work were supported by the NIH R01NS052741-10, NIA R01AG058812, the National Multiple Sclerosis Society (G-1508-05951, RG-1901-33209 and FG-1908-34819) and the Minnesota State Spinal Cord Injury and Traumatic Brain Injury Research Program. 78c-containing chow was generously provided by Calico Life Sciences LLC. Diagrams were created using elements from the Biomedical-PPT-Toolkit-Suite (Motifolio).

## Notes

**Conflict of Interest**: ENC holds a patent on the use of CD38 inhibitors for metabolic diseases. The other authors declare no competing financial interests.

### Competing Interest Statement

ENC holds a patent on the use of CD38 inhibitors for metabolic diseases. The other authors declare no competing financial interests.

