## Supplemental Figure 1 for "Critical role for astrocyte NAD^+^ glycohydrolase in myelin injury and regeneration"

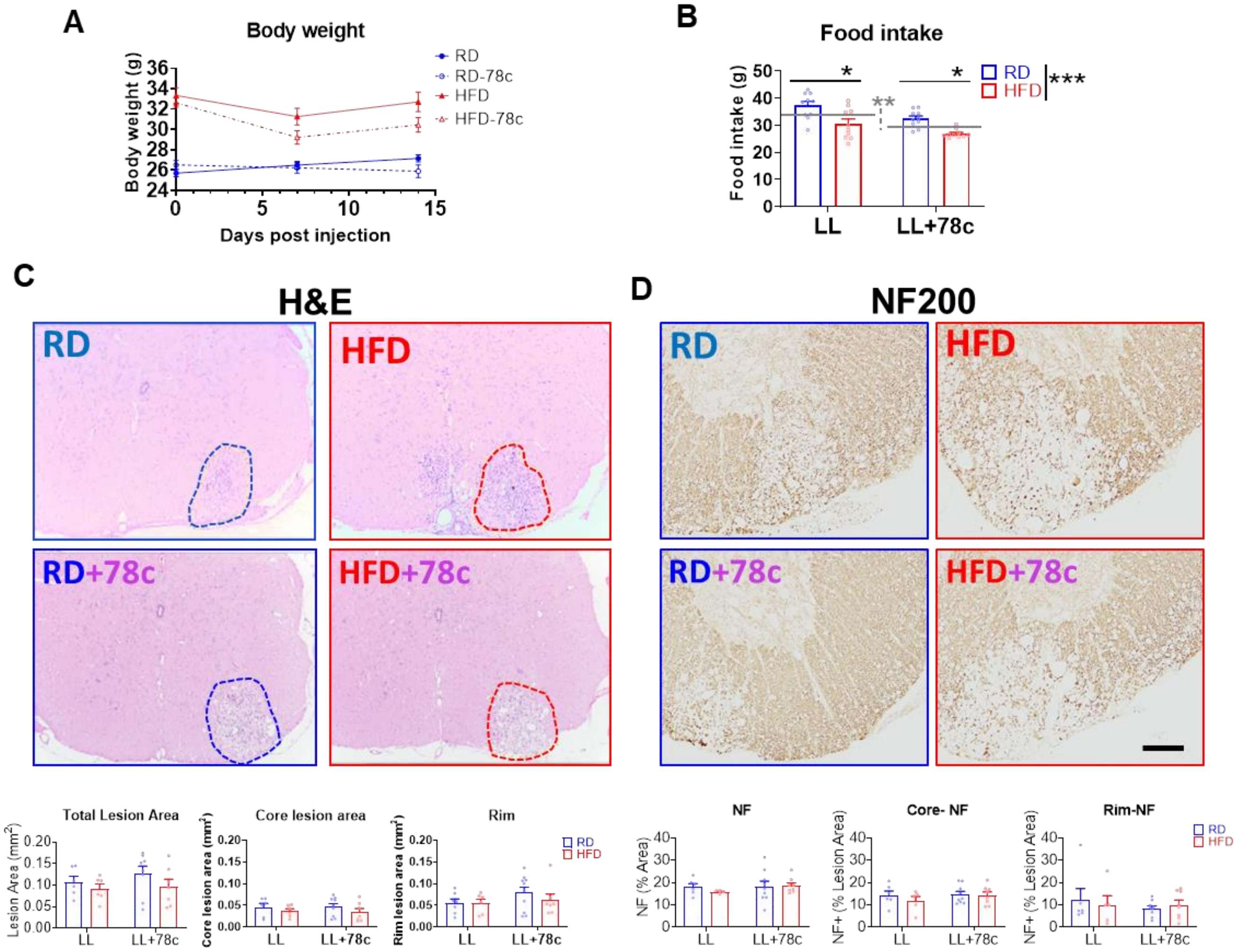


**Supplemental Figure 1. Lysolecithin-induced demyelination model characterization.**

Eight week old male C57Bl6 mice were fed a regular diet (RD) or high fat diet (HFD) for 5 wks, and then split into two additional groups: one receiving the same diet and the other half receiving the respective diet with 78c added (600 ppm). One week later, a focal demyelinating lesion to the ventral spinal cord was induced by lysolecithin (LL, 1% with Evans Blue dye) and mice were allowed to recover for two weeks. Lesion boundary regions were determined by limiting to the area of continuous myelin loss (core) or by including the entire inflammatory lesion (rim) based on immunohistochemical analyses.

**A** Body weights were not significantly affected by 78c, but were by high fat diet (HFD) consumption when compared to regular diet (RD).

**B** Food intake was significantly altered by HFD or 78c consumption.

**C** H&E staining shows the lesion area (C, entire boundary outlined by dotted line and determined by consideration of all stains performed). No significant differences between RD and HFD groups were noted.

**D** Neurofilament, (a marker of axon integrity) as determined by immunohistochemical quantification, was unchanged by either drug or diet.

Bar graphs represent mean +/- SEM. Two-way ANOVA (n=7-10 mice/group). Asterisks indicate levels of statistical significance, where *, p<0.05 and **, p<0.01. Gray line indicates treatment group means,  with gray broken line indicating comparison between treatment group means.
